# Wheat inositol pyrophosphate kinase TaVIH2-3B modulates cell-wall composition for drought tolerance in Arabidopsis

**DOI:** 10.1101/743294

**Authors:** Anuj Shukla, Mandeep Kaur, Swati Kanwar, Gazaldeep Kaur, Shivani Sharma, Shubhra Ganguli, Vandana Kumari, Koushik Mazumder, Pratima Pandey, Hatem Rouached, Vikas Rishi, Rashna Bhandari, Ajay K Pandey

## Abstract

**Background:** Inositol pyrophosphates (PP-InsPs) are high-energy cellular molecules involved in different signalling and regulatory responses. Two distinct classes of inositol phosphate kinases responsible for the synthesis of PP-InsPs have been identified in *Arabidopsis* i.e. ITPKinase (inositol 1,3,4 trisphosphate 5/6 kinase) and PP-IP5Kinase (diphosphoinositol penta*kis*phosphate kinases). Plant PP-IP5Ks are capable of synthesizing InsP_8_ and were shown to control pathogenic defence and phosphate response signals. However, other roles offered by plant PP-IP5Ks, especially towards abiotic stress, remain poorly understood.

**Results:** Here, we characterized two *Triticum aestivum* L. (hexaploid wheat) PPIP5K homologs, *TaVIH1* and *TaVIH2*, for their physiological functions. We demonstrated that wheat VIH proteins could utilize InsP_7_ as the substrate to produce InsP_8_, a process that requires the functional VIH-kinase domains. At the transcriptional level, both *TaVIH1* and *TaVIH2* are expressed in different wheat tissues, including developing grains, but show selective response to abiotic stresses during drought-mimic experiments. Overexpression of *TaVIH2-3B* homolog in *Arabidopsis* conferred tolerance to drought stress and rescued the sensitivity of *Atvih2* mutants. RNAseq analysis of *TaVIH2-3B* transgenic lines of *Arabidopsis* showed a genome-wide reprogramming with remarkable effects on cell-wall biosynthesis genes with enhanced the accumulation of polysaccharides (arabinogalactan, cellulose and arabinoxylan).

**Conclusions:** Overall, this work identifies a novel function of VIH proteins, implying their roles in modulating cell-wall homeostasis genes and providing water-deficit stress tolerance. This work suggests that the plant VIH enzymes could be linked to drought tolerance and also opens up investigations to address the roles of plant VIH derived products in generating drought resistant plants.

## Background

Inositol phosphates (InsPs) are a well-known family of eukaryotic water-soluble signalling molecules that are conserved mainly in their function [1, 2]. This family is characterized by the presence of phosphate either at the single or all the 6-carbon inositol ring backbone. The full phosphorylated InsPs (InsP_6_; *myo* inositol hexakisphosphate, phytic acid) species can be again phosphorylated to generate high energy Inositol pyrophosphates (PP-InsPs)[3–5]. PP-InsPs are essential members of the inositol poly-phosphate family, with an array of pyrophosphates chains present at specific positions [6, 7]. The two major members of InsPs, i.e., InsP_7_ and InsP_8_, are present in very low abundance in cells and are synthesized by two classes of enzymes. The first class of enzyme, inositol hexakisphosphate kinases (IP6Ks), phosphorylate one of the precursors InsP_6_ to form PP-InsP_5_ [3, 8]. The second class of enzyme, diphosphoinositol pentakisphosphate kinases (PP-IP5Ks), phosphorylate InsP_7_ to form InsP_8_ /1,5PP-IP_4_ [5, 9, 10].

During the past two decades, three isoforms of IP6Ks (IP6K1, IP6K2 and IP6K3) and two PP-IP5K (PP-IP5K1 and PP-IP5K2) were identified in humans and mouse [11, 12]. In yeast, a single IP6K (also referred to as Kcs1) and a PP-IP5K (also known as Vip1) are involved in the synthesis of the respective forms of InsP_7_ and InsP_8_ [5, 10]. These high energy pyrophosphates participate in cellular activities such as DNA recombination, vacuolar morphology, cell-wall integrity, gene expression, pseudohyphal growth and phosphate homeostasis as demonstrated in yeast, mice and humans [13–19].

Earlier, the presence of high anionic forms of InsP_6_ was predicted in plant species such as barley and potato [20, 21]. However, the quest to identify the plant genes encoding for these inositol pyrophosphate kinases remained elusive till the identification of two plant VIP genes from *Arabidopsis* and are present in all available plant genomes [5, 17]. In plants, VIP-homolog, also referred to as VIH proteins, contains bifunctional domains including “rim-K” or ATP-grasp superfamily domain at the N-terminal and histidine acid-phosphatase domain at a C-terminus as in yeast [5, 22–24]. Furthermore, these VIH proteins displayed PP-IP5K-like activity involving in plant defence response mediated through jasmonate levels [22].

Recent evidence about genetic interaction studies implies that deletion of the VIH1 and VIH2 in Arabidopsis affects plant growth and is an integral part of the phosphate (Pi) response pathway [23]. The enzymatic properties of *Arabidopsis* VIH1 and VIH2 suggest that both could utilize PP-InsP_5_ as a substrate, akin to the human PP-IP5K2 activity [9, 23]. Additionally, these VIH proteins were functionally active and could rescue invasive growth through hyphae formation in yeast *vip*1Δ mutants [25]. The new line of evidence also suggested that the generated InsP_8_ could bind the eukaryotic SPX domain and thereby regulate the activity of the phosphate starvation response1 (PHR1), a central regulator of phosphate (Pi) starvation [17, 23]. The conserved role of VIH kinases in synthesizing PP-InsPx are essential for their role in Pi homeostasis as demonstrated in yeast, humans and plants [17, 24, 26]. Thus, the role of plant VIH and PP-InsPs need further investigation to explore their additional molecular functions.

In summary, to date, the studies could reveal the function of plant VIH only in pathogen defence and Pi limiting conditions. Still, no other role has been investigated or reported for these genes in *Arabidopsis* or other crop plants. In the current study, we have identified two functionally active VIH genes from hexaploid wheat (*Triticum aestivum* L.), capable of utilizing InsP_7_ as a substrate to generate InsP_8_. Hexaploid wheat, an important crop around the globe and its productivity can be affected when exposed to abiotic stress [27]. We have done expression studies, physiological investigations accompanied by forward and reversed genetic approaches; we provide evidence that wheat VIH2 could impart tolerance to drought in transgenic *Arabidopsis.* Further, we observed that the drought tolerance was dependent upon distinct transcriptomic re-arrangements in addition to alterations in the composition of plant cell-wall. Together, our study provides novel insight into the possible function of plant VIH towards stress tolerance.

## Results

### Phylogeny and Spatio-temporal characterization of VIH genes in wheat tissue

Our efforts to identify potential wheat VIH-like sequences revealed six genes with three homoeolog, which shows 98.8% sequence identity with each other. *TaVIH1* and *TaVIH2* were mapped to chromosome 3 and 4, respectively. Both the wheat VIH genes were present on all the three genome-homoeologs (A, B and D). The Kyte-Doolittle hydropathy plots indicated that wheat VIH proteins were devoid of any transmembrane regions (Supplementary Figure S1A and Table S1). Phylogenetic analysis clustered plant VIH homologs together with TaVIH proteins close to *Oryza sativa* (∼90%) in the monocot specific clade (Figure 1A). TaVIH2 is closer to AtVIH1and AtVIH2 in the phylogenetic tree with an identity of 72% and 33% with ScVIP1, respectively. TaVIH1 reveal a high identity of 78 % with Arabidopsis (AtVIH) proteins and 35% with yeast (ScVIP1) proteins but present in different clad of the tree (Figure 1A). Among themselves, wheat VIH1 (TaVIH-1) and VIH2 (TaVIH-2) show 70% sequence identity at the protein level (Supplementary Figure S2). Amino acid sequence alignment of wheat VIH protein sequences suggested the presence of conserved dual-domain architecture with two distinct domains consisting of N-terminal rimK/ATP GRASP fold and a C-terminal histidine acid-phosphatase (HAP) of PP-IP5K/VIP1 family (Supplementary Figure S1B).

**Figure 1.**
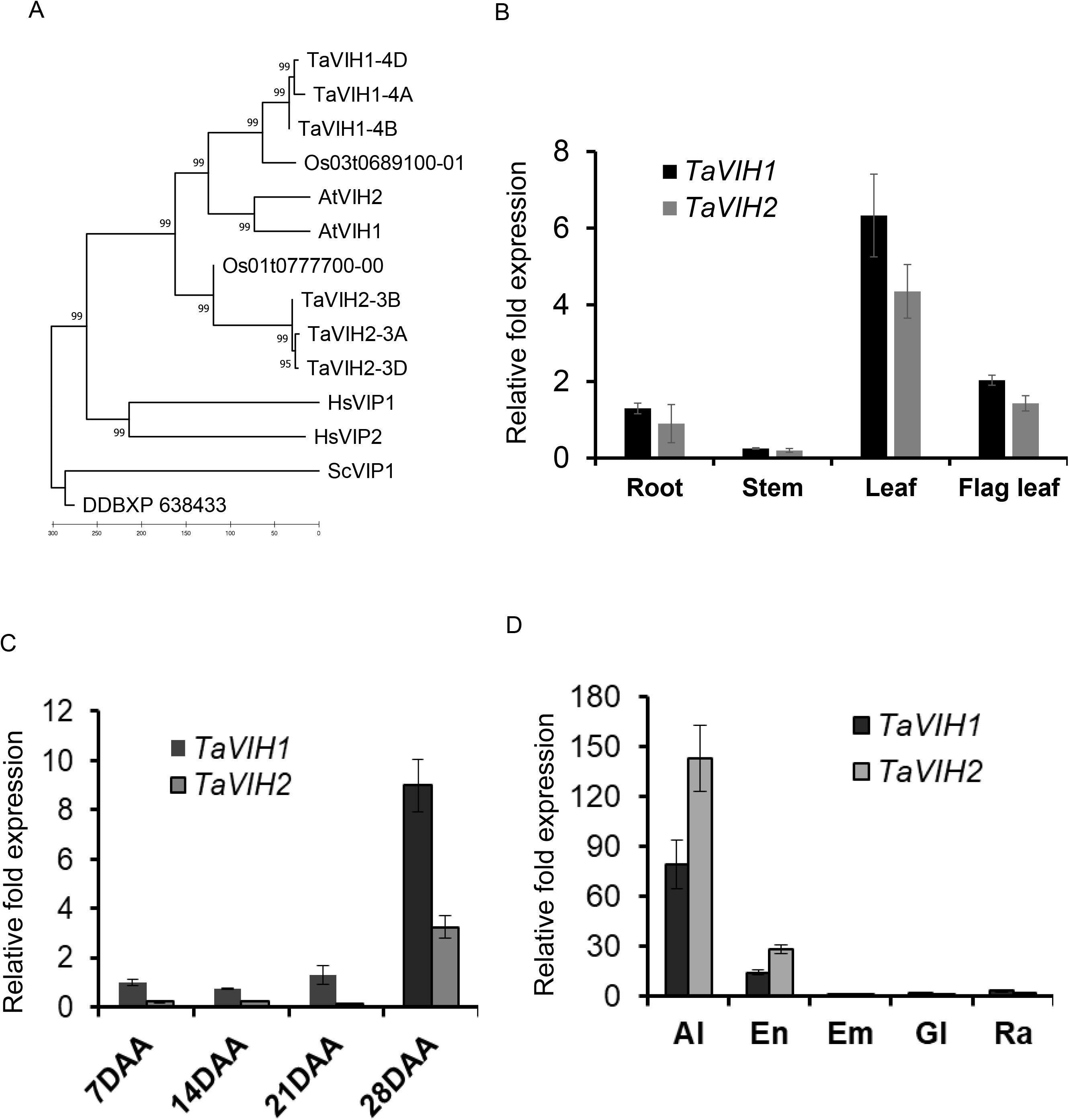
Neighbourhood-Joining phylogenetic tree and expression analysis of wheat genes encoding VIH. (A) Neighbourhood-Joining phylogenetic tree of PP-InsP_5_ proteins. The full-length amino acid sequences of VIH proteins from various taxonomic groups were used for the construction of phylogeny using MEGA7.0. The number represents the bootstraps values (1000 replicates). For construction of evolutionary history was inferred by Minimum Evolution method using 14 amino acids sequences spanning from all the wheat VIH homoeologs (TaVIH1 and TaVIH2), rice (Os01t04777700; Os03t0689100), *Arabidopsis thaliana* (NP_568308; NP_186780), human (HsVIP1-NP_001124330 and HsVIP2-NP_001263206), yeast (VIP1-NP_013514) and *Dictyostelium discoideum* (DDBXP638433) (B) *TaVIH1* and *TaVIH2* in different tissues of a wheat plant. The cDNA was prepared from 2µg of DNA free-RNA isolated from root, stem, leaf and flag leaf tissues of a 14 DAA plant as a template. (C) Quantitative expression analysis of *TaVIH* genes at different seed maturation stages (7, 14, 21 and 28 days after anthesis and; (D) Expression in the tissue of 14 DAA seed (aleurone, Al; endosperm, En; embryo, Em; glumes, Gl and rachis, Ra. For qRT-PCR, cDNA was prepared from 2µg of DNA-free RNA isolated from respective tissues. *TaARF* was used as an internal control for the normalization of Ct values. Standard deviation was calculated along with its level of significance at p<0.05 (*) with respect to the first tissue.

Transcript accumulation of *TaVIH* genes showed similar expression profiles for both genes, with the highest expression in leaf tissues followed by flag leaf, root, and slightest expression in the stem of wheat (Figure 1B). These findings suggest that both VIH genes are preferentially expressed in leaf (Figure 1B). The highest expression of both VIH genes was observed at late stages of grain filling with high transcript accumulation at 28 DAA stage (Figure 1C). Similar levels of transcript accumulation were found in the remaining grain tissues, *viz.* embryo, glumes, and rachis, suggesting a ubiquitous expression in these tissues (Figure 1D). The expression profile in different grain tissues also revealed higher expression of *TaVIH2* genes in the aleurone layer and endosperm tissue which is ∼2-fold higher than TaVIH1 (Figure 1D). Thus, our analysis shows differential expression patterns of VIH in different wheat tissue.

### Wheat inositol pyrophosphate kinase demonstrates PP-IP_5_K activity

Yeast complementation assay for wheat VIH genes was carried out using yeast growth assay on SD-Ura plates supplemented with 0, 2.5 and 5 mM 6-azuracil. The expression of both TaVIH1-4D and TaVIH2-3B in yeast was confirmed by western blotting (Supplementary Figure S3A). The wild type strain BY4741 showed an unrestricted growth phenotype, whereas *vip1*Δ transformed with empty pYES2 vector showed growth sensitivity at 2.5 and 5mM concentrations of 6-azauracil [22] (Supplementary Figure S3A). To our surprise, the mutant strain transformed with pYES2-TaVIH1-4D could not revive growth defect on selection plates, whereas the pYES2-TaVIH2-3B could rescue the growth phenotype of the *vip1*Δ strain. Previous studies show that under stress conditions, unlike wild type yeast, *vip1*Δ mutant does not form pseudo-hyphae [25]. The complemented *vip1*Δ strain with pYES2-TaVIH2-3B could also rescue this phenotype during stress by showing hyphae formation (Supplementary Figure S3B). Overall, our data suggest that *TaVIH2* derived from the B genome can complement the growth defects of the *vip1*Δ strain

In-vitro kinase assay was performed using the pure protein of VIH-kinase domain (KD) (Supplementary Figure S4). Firstly, we generated InsP_7_ substrate using mouse IP6K1 enzyme using InsP_6_. The synthesized InsP_7_ was confirmed by TBE-PAGE gel (Supplementary Figure S5A), and the gel eluted product was also subjected to MALDI-ToF (Supplementary Figure S5B). The relative luminescence units (RLU) were recorded for TaVIH1-KD and TaVIH2-KD using mIP6K1 generated InsP_7_ as a substrate (Figure 2A). The RLU value represents the ADP formed during the kinase reaction. Our assays show a significant increase in the RLU for both the TaVIH proteins in the presence of InsP_7_ substrate. Among them, the wheat VIH2 showed a high fold luminescence response compared to the VIH1 protein (Figure 2A). This kinase activity was diminished in VIH post-heat-denaturation (D-VIH), and the activity was not significantly different when compared to either Enzyme-control (Ec) or substrate control (InsP_7_) reactions. This conversion of ATP to ADP can be used as an indirect measurement biosynthesis of InsP_8_.

**Figure 2:**
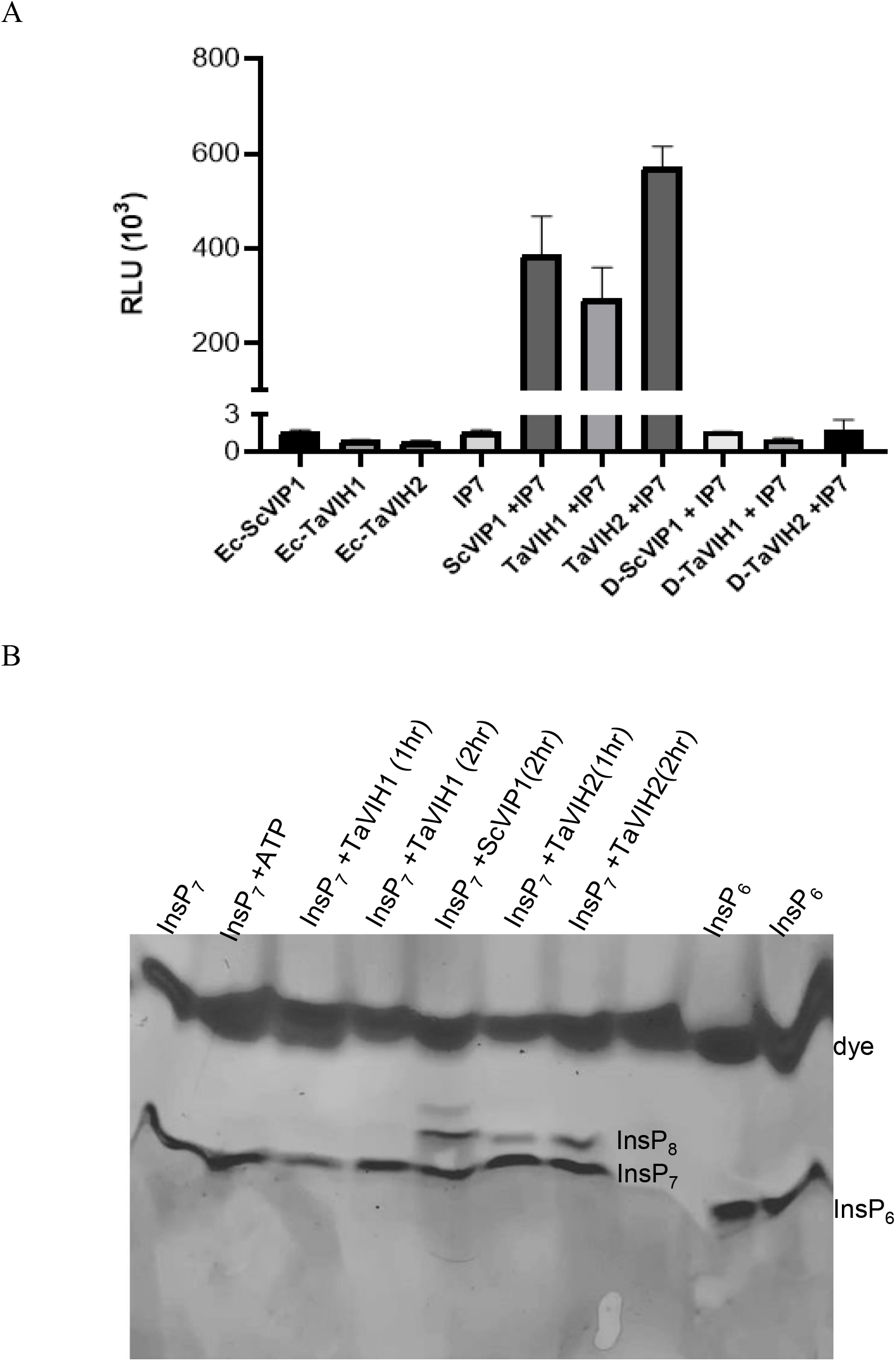
Enzymatic activity and analysis of the PP-InsP on PAGE. (A) The relative luminescence units for all reactions performed were recorded using Spectramax optical reader. The kinase reactions were performed using 50 ng of TaVIH1-KD and TaVIH2-KD purified proteins for 30 mins, followed by steps mentioned in the ADP-GLO kit. (B) Visualization of PP-InsP products on the PAGE-GEL (33%). The In-vitro kinase reactions were performed using 30 ng of ScVIP1-KD, TaVIH1-KD and TaVIH2-KD purified proteins for 1 and 2 hr at 28° C. The reactions were then resolved on the gels (TBE-PAGE). The photo was taken after staining by Toluidine Blue.

The InsP_8_ product generated by the above reactions was confirmed by resolving the reaction products by TBE-PAGE analysis [28]. To visualize the products on a gel, we used a higher concentration of InsP_7_ substrate. As a control, we used ScVIP1-KD generated InsP_8_ using InsP_7_ as a substrate (Figure 2B; lane5). The TaVIH proteins were incubated with InsP_7_ as a substrate for two-time points (1 and 2 hr), and the products were resolved by PAGE. Our experiments suggest that InsP_8_ was synthesized only by TaVIH2-KD when InsP_7_ was provided as a substrate (Figure 2B). During this period of incubation, no detectable levels of the product were seen for the TaVIH1-KD reactions. In contrast, upon a longer incubation with substrates (∼9 hrs), we observed that InsP_8_ was generated by both VIH1 and VIH2 proteins (Supplementary Figure 5C), suggesting that TaVIH1 may have a lower enzyme activity compared with TaVIH2. To further confirm the nature of generated phosphorylated inositol molecules, MALDI-ToF-MS was performed. The analysis of the InsP_8_ band (generated by TaVIH2-KD) was done in the m/z range of 500 to 1000, which reveals a significant peak of 820.47 m/z (Supplementary Figure 5D). Here minor peak represents the theoretical mass of InsP_8_ and the prominent peak corresponding to the InsP_8_-acetonitrile adduct. These enzymatic and analytical experiments confirm that TaVIH2 protein is functionally active and capable of using InsP_7_ as a substrate under in-vitro conditions and may possess PP-IP_5_K like activity.

### Expression of *35S: TaVIH2-3B* transgenic *Arabidopsis* display robust growth

The biological functions of TaVIH2 were analyzed by overexpressing the cDNA of TaVIH2-3B in Columbia (Col-0) *Arabidopsis thaliana*. In total, seven transgenic lines were pre-selected based on TaVIH2 expression that was analyzed by western analysis (Figure 3A). Further, four transgenic lines (#Line2-3, #Line 4-3, #Line 5-2 and #Line 6-1) were selected for characterization. We observed that at the vegetative stage, TaVIH2-3B transgenic *Arabidopsis* showed robust growth. Plants (14 days old seedlings) showed enhanced rosette area cover and increased number of leaves as compared to the controls (Col-0 and Col-0Ev (empty vector))(Figure 3B, C and D). These transgenic *Arabidopsis* also displayed enhanced branching with an overall increase in the length of the main shoot axis and leaf size as compared to the controls (Figure 4A and B). Primary and secondary shoot numbers were also enhanced in the transgenic *Arabidopsis* (Figure 4D). In general, no significant differences during the flowering stage was observed, yet the increased number of (20-24) secondary shoots were evident when compared with control plants (12-15 shoots) (Figure 4D and E). These results suggest that the expression of TaVIH2-3B in *Arabidopsis* impacts the overall growth of the plant.

**Figure 3:**
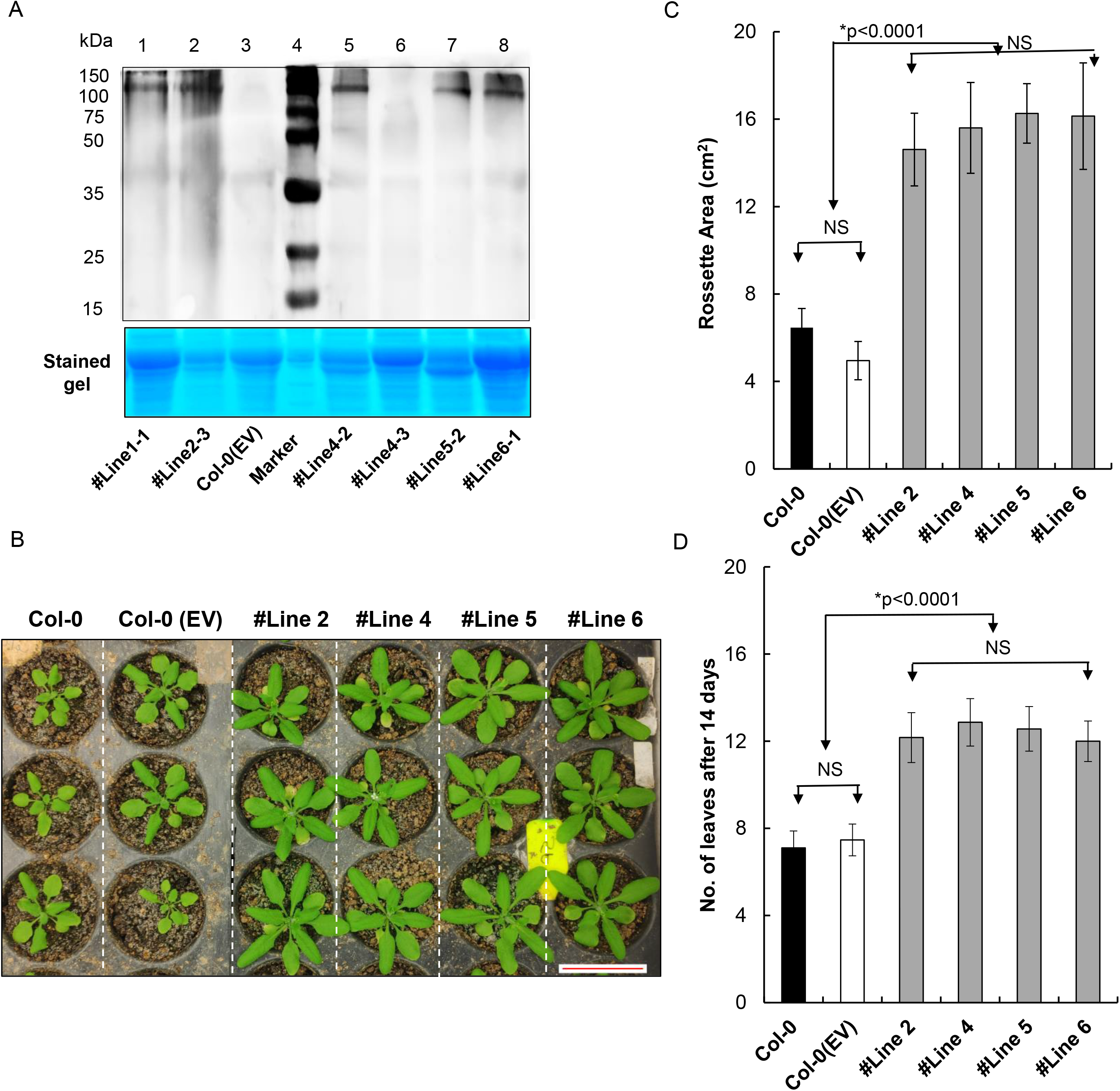
Generation of VIH2-3B transgenic Arabidopsis and its characterization. (A) Western analysis and screening of Col-0 Arabidopsis transgenic lines for TaVIH2-3B protein (∼100 kDa) overexpressing lines. Multiple transgenic lines were screened, and Western was done using His-Antibody using 20µg of total protein. Coomassie Blue stain of the total protein (lower panel) was used as a loading control. (B) Representative picture of the rosette area of the Transgenic Arabidopsis (#Line2, #Line4, #Line5 and #Line6) and controls. (C) Rosette area measurement (in cm2) using Image-J for 4 different transgenic lines along with the controls. Measurement was taken after 14 days of growth. (D) Number of Rosette leaves in transgenic Arabidopsis and control lines. Three experimental replicates using 10 plants each were used to calculate the parameters.

**Figure 4:**
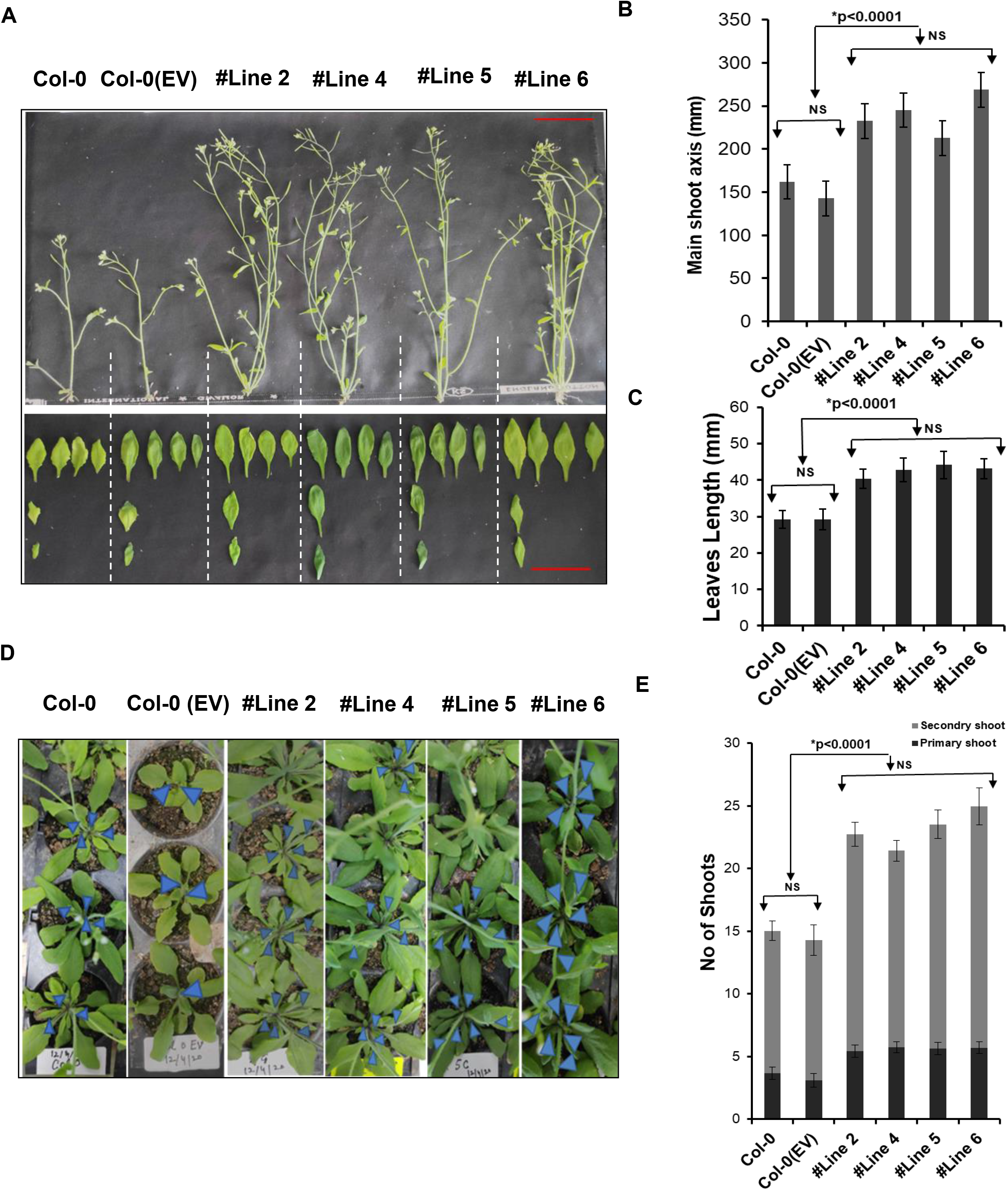
Morphological characterization of VIH2-3B transgenic Arabidopsis. (A) representative phenotype of transgenic Arabidopsis and controls post 25 days of growth (flowering stage). (B) length of the main axis. (C) leaves size (in mm). (D) Phenotype for the shoots of the transgenic Arabidopsis and controls. The panel indicates the morphological differences in the number of primary shoots (as indicated by arrows) among the lines. (E) A number of total shoots (primary and secondary shoots) in transgenic Arabidopsis and control plants (right panel). For each transgenic line, three experimental replicates were performed using 10 plants each.

### Wheat VIH2-3B respond to drought-mimic stress

To investigate the promoter activities of *TaVIH1* and *TaVIH2*, 5’ flanking regions (1 kb) of these genes were cloned, and the comparative analysis revealed the presence of hormones and abiotic stress-responsive cis-elements (Supplementary Figure S6A). The presence of these elements suggested that wheat *VIH* could be regulated by stress. Notably, we observed the presence of the cis-elements that could respond to drought/dehydration, P1BS (PHR1 binding site) and GA responsive domains. (Supplementary Figure S6A). This motivated us to perform preliminary screening experiments using *TaVIH*-promoters fused to β-glucuronidase (GUS)-reporter gene (pVIH1/2:GUS) in *Arabidopsis* (Col-0). A significant increase in GUS reporter activity of *pVIH2:GUS* lines indicated the ability of this promoter to sense the given stress and drive GUS reporter expression. Interestingly, the *TaVIH2* promoter responded strongly to dehydration/drought stress and Pi-starvation (Supplementary Figure S6B). Subsequently, the GUS was expressed strongly during the presence of 30% PEG (Figure 5A). This suggests the potential role of *TaVIH2* during the drought response. A weak expression of the *TaVIH2* promoter was observed in the presence of ABA and GA_3_ (Supplementary Figure S6B). Control (EV) seedlings showed no visible GUS staining. Based on our reporter assays, we speculate that *TaVIH2* could have an essential role during a drought stress response, which was investigated further.

**Figure 5:**
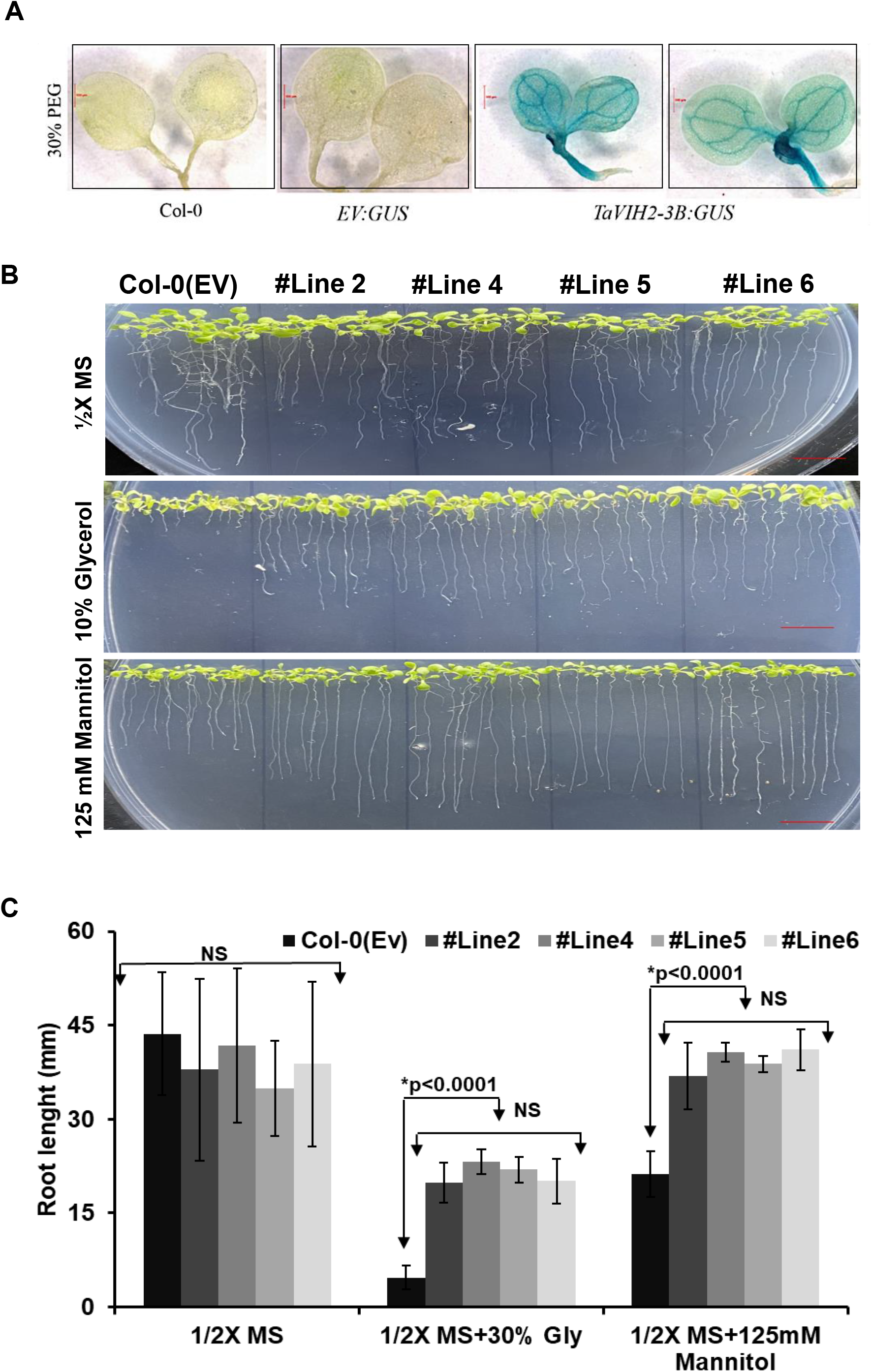
Drought-mimic stress for VIH2-3B *Arabidopsis* transgenic lines. (A) Reporter assays using promTaVIH2:GUS transgenic lines subjected to drought-mimic (30% PEG). Seedlings with or without treatment (control) were stained overnight in GUS staining solution and photographed using Leica stereomicroscope at 6.3X magnification. (B) Transgenic Arabidopsis and control seedlings were subjected to drought mimic conditions with glycerol-10 % and mannitol-125 mM. Ten seedlings were used for each transgenic line for each treatment. These experiments were repeated in three experimental replicates with a similar phenotype. (C) Root length of treated seedlings (in mm) for all the lines. Twenty seedlings were used for the measurement of root length for each line.

We tested the gene response to drought-like conditions on plant physiology. Here, seedlings were exposed to drought-like conditions using mannitol (125 mM) and glycerol (10 %) [29]. No significant difference in the root growth pattern on the ½ MS plates was observed in all the *Arabidopsis* seedlings (Figure 5B). Inhibition of the root growth was observed for the control *Arabidopsis* suggesting their sensitivity to the presence of both the mannitol and glycerol (Figure 5B). In contrast, TaVIH2-3B overexpression in *Arabidopsis* was able to escape the detrimental root growth (Figure 5C). Finally, to check the sensitivity of *Arabidopsis vih2-3/vih2-4* and its rescue by wheat VIH2, we screened the complemented mutant lines with *TaVIH2-3B* and evaluated it during drought-mimic conditions (Figure 6A). Interestingly, both *vih2-3* and *vih2-4* showed high sensitivity towards drought mimicking conditions and this sensitivity was restored to similar to Col0-(EV) when complemented with *TaVIH2-3B* (Figure 6B and C). These results corroborate the intriguing aspects of TaVIH2 physiological function during drought stress.

**Figure 6:**
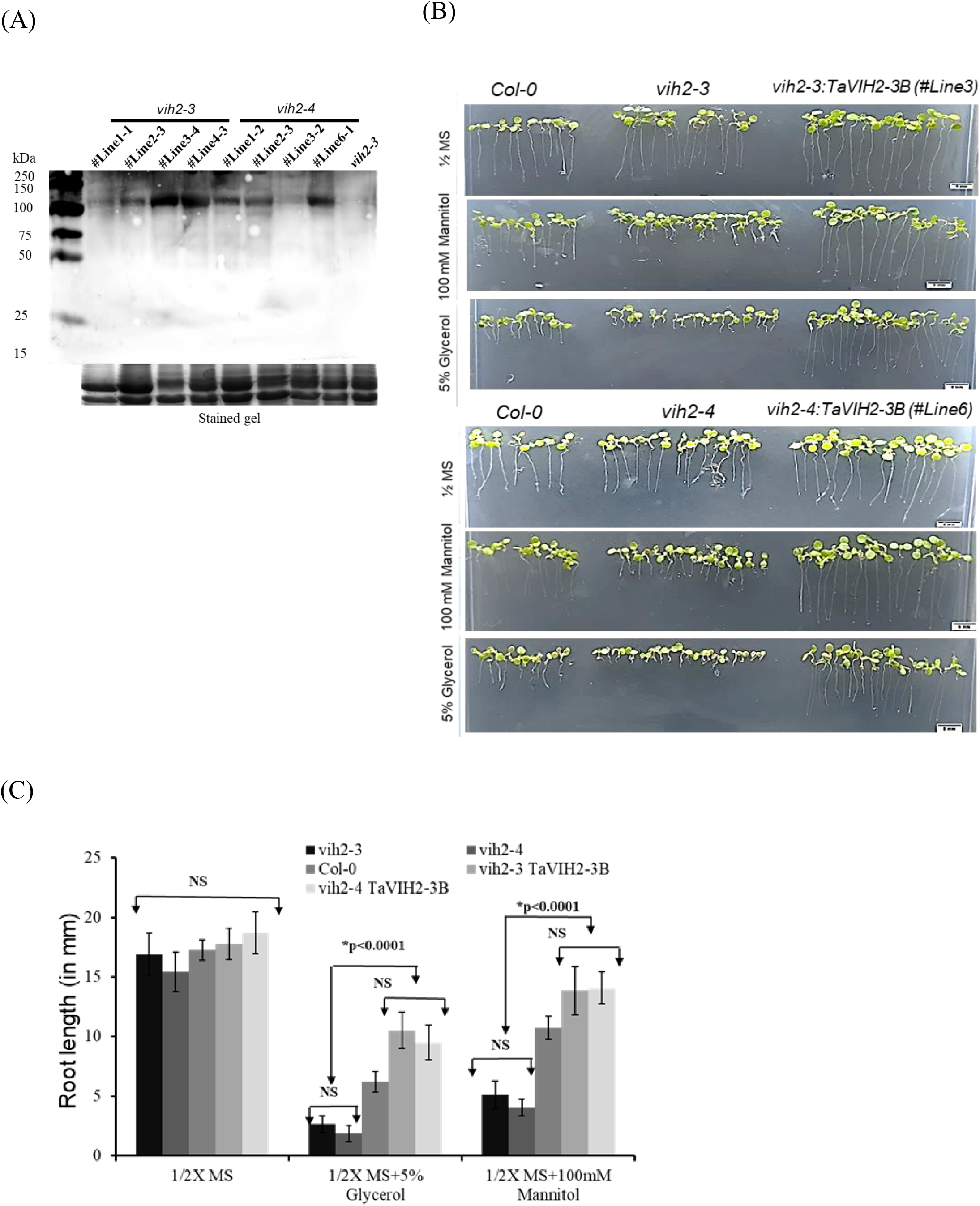
Complementation of *vih2-3* line and its characterization. (A) Western analysis of *vih2* mutant lines (*vih2-3* and *vih2-4*) complemented with *TaVIH2-3B*. Multiple transgenic lines were screened and Western was done using His-Antibody using 20 µg of total protein. Coomassie Blue stain of the total protein (lower panel) was used as a loading control. (B) Transgenic *Arabidopsis, mutant* and Col-0 seedlings were subjected to drought mimic conditions with glycerol-5 % and Mannitol-100 mM (for moderate mimic-drought). Eight to ten seedlings were used for each transgenic lines for each treatment. (C) Root length in mm (n=20). These experiments were repeated three experimental replicates with similar phenotype.

### Wheat VIH2-3B imparts resistance to water-deficit stress

Studying the relative water loss helped us investigate the direct involvement of TaVIH2-3B in conferring the drought tolerance in the detached leaves. The rate of water loss was very significant in the control plants compared to the transgenic plants (Figure 7A). This loss was less in the transgenic *Arabidopsis* (40-46%) when compared to control plants (16-18%) after 8 hrs of incubation (Figure 7A). Next, we measured relative leaf water content (RWC%; Figure 7B) for these plants. The RWC was high (∼65%) in transgenic plants as compared to the control plants (∼46%).

**Figure 7:**
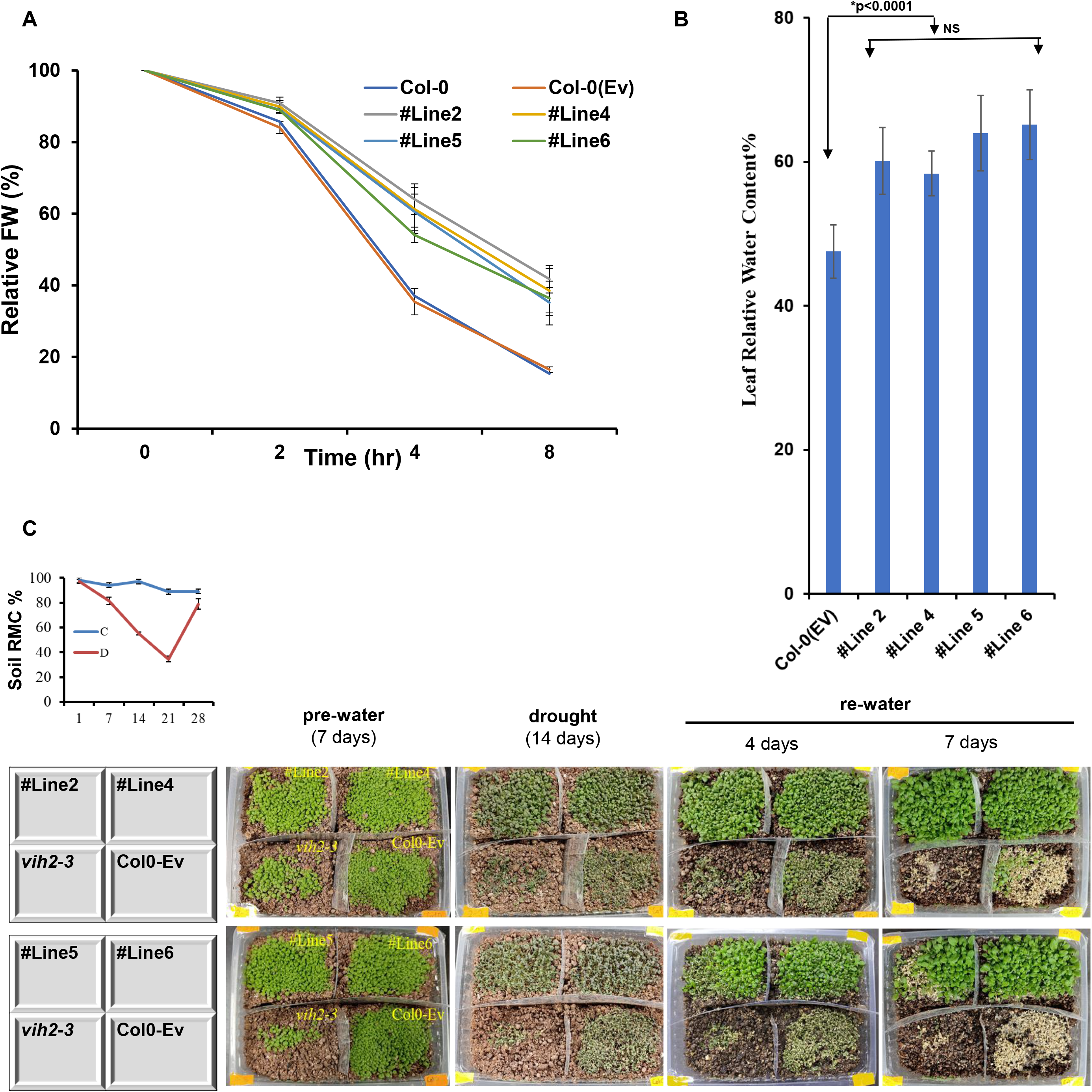
Drought response of VIH2-3B *Arabidopsis* transgenic lines. (A) Relative water loss in Arabidopsis leaves post 8 hrs. Three experimental replicates, each with ten leaves, were used to calculate the water loss %. (B) Leaf relative water content was measured after 24 days of growth. (C) Drought treatment of soil-grown plants. Seedlings were pre-grown for the period of fourteen days were subjected to drought for additional fourteen days. The plants were then re-watered for a period of seven days, and % survival rates were calculated. This experiment was repeated twice. Representative pictures were taken post seven days of re-watering.

Further, drought stress experiments were carried out for all the plants by exposing 7 days old plants to 14 days of water withholding (drought). These experiments were carried out for the mutant, wild-type and overexpressing plants together in the same pot for ensuring that they are inter-rooted and exposed to the same soil moisture conditions. After 14 days of drought, the relative soil moisture content was observed to be as low as 35% in the pots. This caused a dramatic withering of both control and transgenic *Arabidopsis* plants. However, when the plants were re-watered, high survival rates (∼65 %) were observed in the transgenic plants, whereas no or very low (3%) survival efficiency was observed in control. No survival was observed for the *vih2-3* mutant plants (Figure 7C), indicating their sensitivity to drought conditions. This indicates that the transgenic *Arabidopsis* overexpressing TaVIH2 escapes the effect of drought and improves survival rate by imparting drought tolerance.

### Transcriptomics data suggest that *VIH2-3B* stimulate genes related to drought stress

In order to understand the basis of robust phenotype and drought resistance observed in the transgenic Arabidopsis plants when complemented with TaVIH2-3B, we used the transcriptomics approach. Transcriptomics changes in 25 days old seedlings of control and two transgenic plants (#Line4 and #Line6) were analyzed. PCA of normalized expression abundances revealed a high level of correlation among biological replicates (n=3) in each transgenic line. PCA also indicates a distinct cluster for overexpressing transgenic lines and controls (Supplementary Figure S7A). Based on an analysis involving respective three biological replicates, a total of 626 and 261 genes were significantly up and down-regulated (−1>Log FC >1.0) in #Line4 while 797 and 273 genes were up and downregulated in #Line6 transgenic Arabidopsis lines compared to control plants (Supplementary Table S2). Overall, 605 genes were commonly differentially altered in the two transgenic lines with respect to the control plants (Col-0(Ev); Figure 8A).

**Figure 8:**
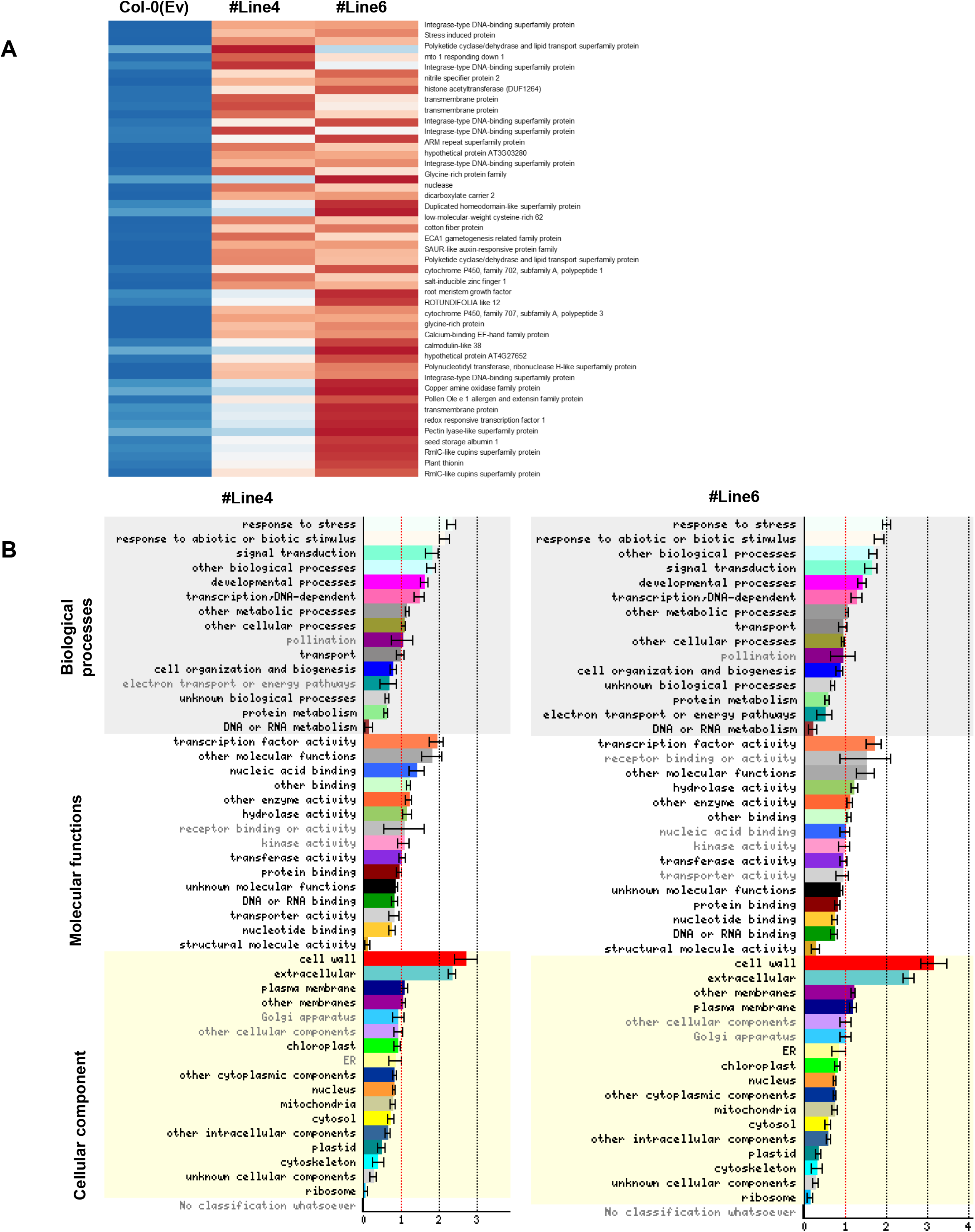
RNAseq analysis of Col-0 and #Line4 and 6. (A) Expression pattern (as Z-scores) of top 56 genes commonly up-regulated among the transgenic lines w.r.t. Col-0(Ev) in 25 days old seedlings. Heatmap depicts the relative expression in Col-0(Ev) and over-expressing lines of TaVIH2-3B (3 biological replicates; rep1-3). (B) Heatmap representing a graphical summary of the Gene Ontology (GO) classification for DEGs in #Line4 and #Line6 w.r.t. Control plants. Increasing intensities of brown and blue colours represent the comparatively low and high expression for each gene, as depicted by the colour scale. Normalized expression counts were used to plotting the expression as Z-scores using heatmap. Two functions from the gplots package in R. Significantly altered GO terms were identified using the Classification SuperViewer tool; the x-axis represents the GO terms where bold terms represent significant alteration while the y-axis represents the normed frequency which when > 1 signifies over-representation while <1 signifies under-representation.

Interestingly, a high number of genes constitutively activated in the transgenic *Arabidopsis* belong to the dehydration response element-binding (DREB) protein, including Integrase-type DNA-binding superfamily proteins and glycine-rich proteins. Upon analysis of the GO terms, the highest number of genes for “stress-related” and “cell-wall related activities” were enriched in the biological process and cellular component categories (Figure 8B and Supplementary Figure S7B). Strikingly, multiple genes involved in cell-wall biosynthesis, modification and degradation were also up-regulated in the transgenic plants (Figure 9A). In addition to that, distinct clusters of genes involved in Abscisic acid (ABA) biosynthesis were also significantly up-regulated among the different lines of transgenic *Arabidopsis* (Figure 9B). Notably, drought-marker genes encoding 9-*cis*-epoxycarotenoid dioxygenase (*AtNCED6* and *AtNCED9*) involved in ABA biosynthesis were also up-regulated. Multiple DREB encoding genes and cytochrome P450 (CYPs) related family genes (*CYP71A23, CYP94B3, CYP71B12, CYP96A2, CYP702A1, CYP707A3, CYP82C2, CYP76G1, CYP705A4, CYP71B10, CYP706A2, CYP81D11*) were also differentially regulated in the transgenic *Arabidopsis* (Figure 9C and D). The expression response of these genes was also validated by using qRT-PCR analysis. Our expression data strongly supported the transcriptome observation that reflects the upregulation of multiple genes (Supplementary Figure S8). These genes validate the abundance of transcripts encoding for DREB, ABA biosynthesis and CYP sub-family genes in transgenic lines when compared to wild type. Overall, we conclude that a distinct cluster of genes involved in drought and ABA stress were significantly up-regulated in these transgenic plants and thus may impart tolerance to stress.

**Figure 9:**
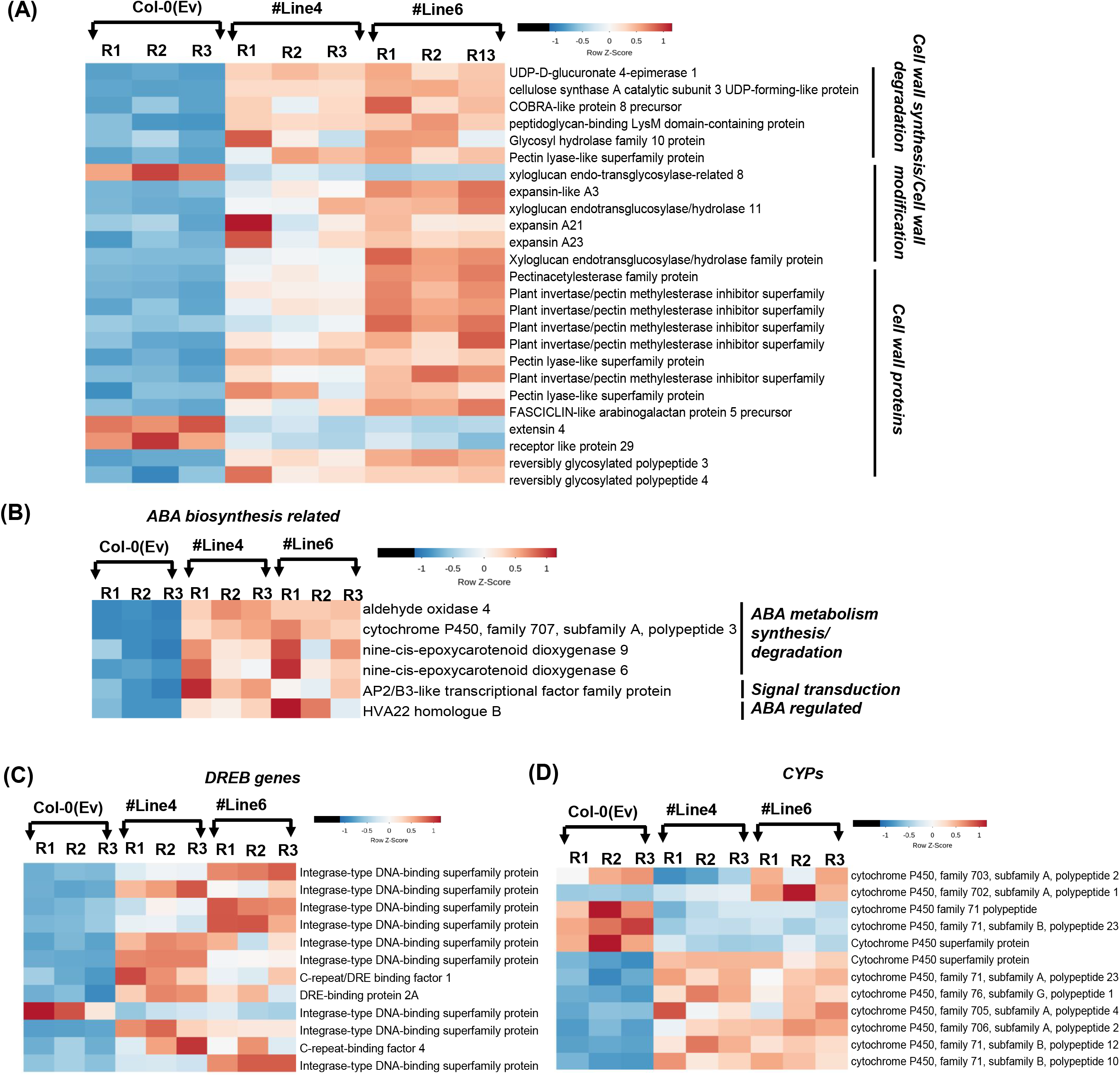
Heatmap expression analysis of gene families in Col-0(Ev) overexpressing *TaVIH2-3B* Arabidopsis (#Line4 and 6). (A) Heatmaps for expression patterns (as Z-scores) for genes DE in both transgenic lines w.r.t. Col-0(Ev), encoding for the genes involved in cell-wall homeostasis. (B) ABA biosynthesis-related pathway genes. (C) DREB encoding genes. (D) cytochrome P450 (CYPs) genes. Increasing intensities of brown and blue colours represent the comparatively low and high expression for each gene, as depicted by the colour scale. Normalized expression counts were used to plotting the expression as Z-scores using heatmap. Two functions from the gplots package (Warnes et al., 2005) in R. Genes encoding for respective pathways were extracted using MapMan (Thimm et al., 2004). R1, R2 and R3 represent the biological replicates for the RNAseq analysis of the individual lines.

### VIH2 overexpression affects ABA levels and regulates plant cell-wall composition

Multiple genes related to ABA biosynthesis were differentially expressed in TaVIH2-3B overexpressing *Arabidopsis*. To verify if the de-novo gene expression response to ABA associated genes could be correlated with its in-vivo levels, ABA was quantified in their leaves. We observed that the accumulation of ABA was significantly higher (∼3-4 fold) in transgenic *Arabidopsis* when compared to the control plants (Figure 10A). This average increase of ABA in all the four transgenic lines was statistically significant (p<0.0001, Student’s t-test). Our data confirmed the involvement of ABA in the drought tolerance of transgenic lines.

**Figure 10:**
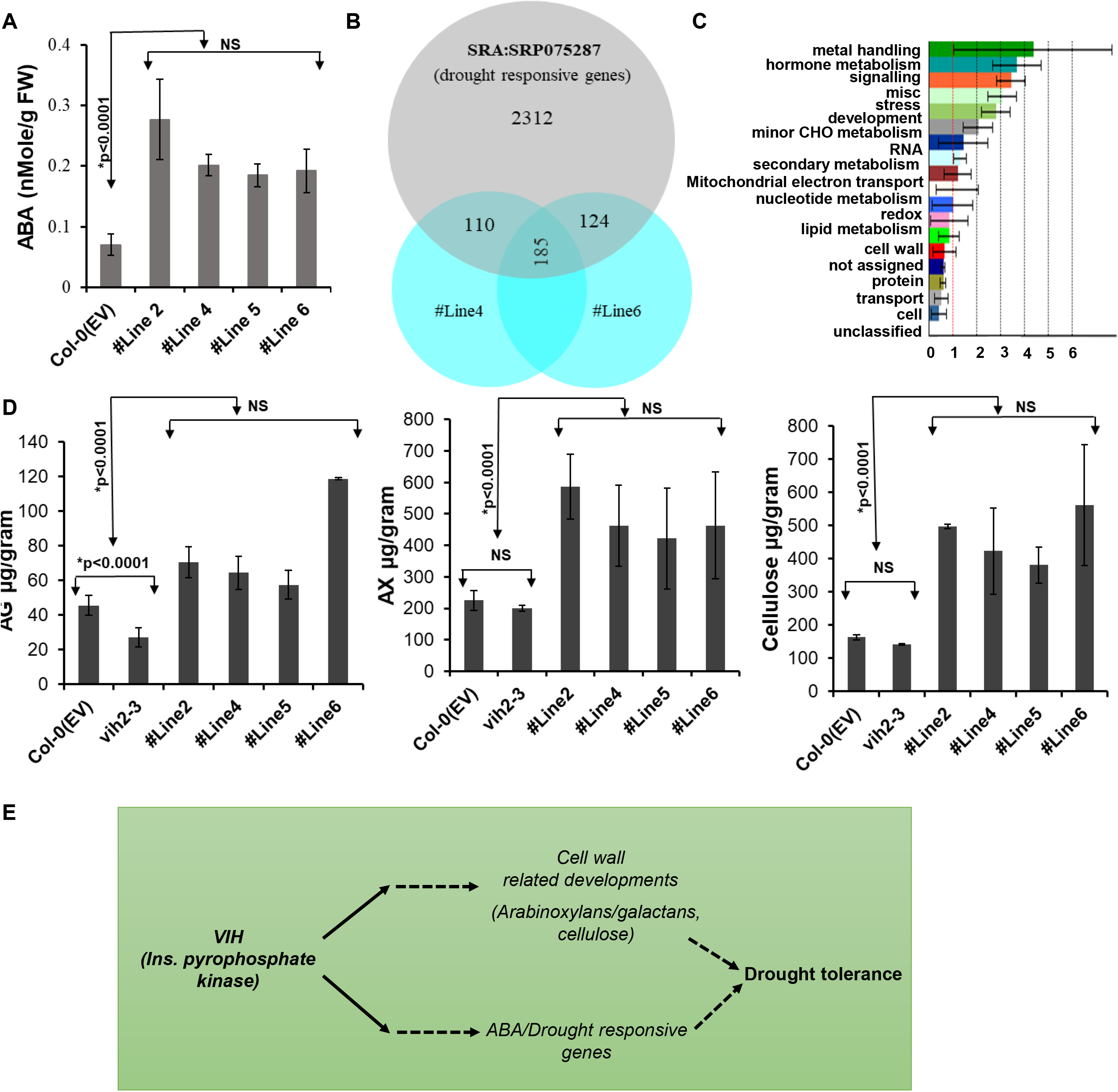
ABA and polysaccharides composition of Arabidopsis shoots. (A) ABA measurement in the leaves of transgenic Arabidopsis overexpressing VIH2-3B and control plants. (B) Venn diagram representation for the genes differentially regulated by drought stress, and transgenic lines #Line4 and #Line6 w.r.t. respective Controls. Drought responsive genes were shortlisted using the Cufflinks pipeline after processing the datasets for 10 days of drought stress and control. (C) Mapman pathway analysis using Classification SuperViewer for the genes that are commonly regulated by drought stress (SRA: SRP075287) as well as transgenic lines w.r.t. control plants (#Line4 and 6). Bold terms represent significant pathways; normed frequency > 1 signifies over-representation while < 1 signifies under-representation. (D) For cell-wall composition analysis, wild-type Col-0, *Arabidopsis* overexpressing TaVIH2-3B (Line#2, 4, 5 and 6) and *Arabidopsis vih2-3* representing Arabidopsis mutant defective for the expression of AtVIH2 were used. Total AG: arabinogalactan, AX: arabinoxylan and cellulose (in µg/g) were measured as indicated in Methods. Analyses were made in triplicates, with each experimental replicate representing at least five plants for each genotype. Vertical bars represent the standard deviation. * on the bar indicates that the mean is significantly different at p < 0.001 (#at p<0.05) with respect to their respective control plants. (E) Speculative model for the working of VIH2 to impart drought resistance to plants

Our results have revealed the function of TaVIH2-3B in drought stress. To draw the commonality between our gene expression in TaVIH2-3B overexpressing *Arabidopsis* and drought, we analyzed previously reported RNAseq data SRA: SRP075287 (under drought stress) for overlap of de-regulated genes. In total, 295 and 309 genes were commonly regulated in #Line4 and #Line6 when compared with drought data (Figure 10B and Supplementary Table S3). Most of the listed genes that were commonly regulated belong to the category of hormone metabolism, signalling, stress response, development and cell-wall functions (Figure 10C). Multiple genes NCEDs, CYPs and glycosyltransferases were highly enriched in the dataset (Supplementary Table S3). These extended analyses support the notion that TaVIH2-3B could impart activation of genes pertaining to drought in transgenic plants that could impart basal drought resistance. Since cell-wall plays a significant role in imparting drought resistance, we, therefore, measured different cell-wall components of control and transgenic *Arabidopsis*. Using standard extraction methods resulted in comparable yields from all the tested plants, and the presence of starch was ruled out before performing experiments. Our extraction procedures for control plants show the ratio of 1::1.2 to 1.5 for arabinose/galactose and arabinose/xylans. This validates our extraction procedures. Our analysis indicated a consistent increase in the accumulation of cellulose (from 1.3 to 2.5-fold) in the transgenic lines that were the same among the biological replicates and multiple transgenic lines (Figure 10D). Additionally, arabinoxylan (AX) and arabinogalactan (AG) was also increased (1.8-2.2 and 1.47-1.5-fold) in the transgenic lines as compared to the controls (Figure 10D). To further validate the role of VIH proteins, the *Atvih2-3* mutant line was used for measuring the biochemical composition of the shoots cell-wall (Figure 10D). Our analysis showed a significant reduction of the AG, AX and cellulose content in this mutant line when compared to transgenic lines (Figure 10D). Our data demonstrate that overexpression of wheat VIH2-3B resulted in changes in the cell-wall composition, and these changes could be linked to the enhanced drought response in leaves.

## Discussion

Recently, studies investigating inositol pyrophosphates have gained much attention due to the presence of high energy pyrophosphate moieties speculated to regulate metabolic homeostasis in organisms [22, 25, 30–32]. This study was performed to characterize and identify the functional mechanism of VIH proteins involved in the biosynthesis of PP-InsPx. We have characterized two wheat inositol pyrophosphate kinase (TaVIH1 and TaVIH2) encoding genes and demonstrated that homoeolog wheat *VIH2-3B* in *Arabidopsis* could enhance growth and provide tolerance to drought stress. Our line of evidence shows that this tolerance to drought is a result of the ability of VIH to modulate cell-wall and ABA related genes resulting in the changes in the cell-wall polysaccharide composition (AG, AX and cellulose).

Hexaploid bread wheat has one of the most complex genomes comprising of three related sub-genomes that have originated from three separate diploid ancestors, thus forming an allohexaploid genome [33, 34]. Therefore, to consider the appropriate homoeolog-transcript for further studies, the Wheat-Exp expression database was used to analyze VIH2-3B homoeolog expression in different tissues and also during the developmental time course (Supplementary Figure S9A). Plant VIHs are known to be involved in defence response via a jasmonate-dependent resistance in Arabidopsis [22]. Wheat VIH genes were also induced upon infection of plants with pathogens (Supplementary Figure S9B and C). Thus, the role of plant VIH genes during plant-microbe interaction was found to be conserved. TaVIH protein was an authentic kinase protein since its kinase domain could catalyze phosphorylation and harbours yeast VIP1-like activity, as demonstrated by its utilization of InsP_7_ as a substrate. In the past, AtVIH proteins possess kinase activity that generates different isoforms of InsP_7_ [22, 25]. Earlier, it was suggested that *Arabidopsis* VIH2 executes Vip1/PP-IP5K but not Kcs1/IP6K-like activities in yeast [22]. This observation confirms the conserved kinase activity among the plants with high substrate affinity for InsP_7_ [35]. Similarly, yeast and human enzymes also show differential InsP_6_ and InsP_7_ kinase activity[5, 36, 37]. We tested InsP_7_ as a substrate for wheat VIH proteins where TaVIH2 shows more specificity towards InsP_7_ that suggest PP-IP5K like activity generating InsP_8_ (Figure 2B). Intriguingly, our time-dependent assays and the RLU value, which reflects the conversion of InsP_7_ to InsP_8,_ could account for the different affinity of wheat VIH proteins (Figure 2A and B). Interestingly, AtVIH1 and AtVIH2 show a high identity (89.8 %) at the protein level whereas, specifically TaVIH1-4D and TaVIH2-3B arising from two different chromosomes show 72 % identity. VIH protein alignment of Arabidopsis and wheat suggest the presence of the conserved residues required for protein-substrate (5-InsP_7_) interactions (Supplementary Figure S2). Although, the conserved-catalytic residues remain same in both the wheat VIH proteins, we could still see changes in the protein sequences in the N-terminal ATP-grasp domains. Wheat genome encodes a total of six VIH proteins that remains to be tested if they could vary in the affinity to utilize the respective substrates. These apparent differences could be intriguing that requires further biochemical investigations.

The presence of various *cis*-acting elements in the promoter region plays an essential role in the transcriptional regulation of genes in response to multiple environmental factors. Our transcriptional activity of *TaVIH2-3B* promoter and expression analysis links TaVIH2-3B with Pi-starvation response (Supplementary Figure S2). This function of inositol pyrophosphate kinases in the regulation of Pi homeostasis seems to be evolutionarily conserved [31, 37]. In Arabidopsis, it was recently demonstrated that VIH derived InsP8 is required to sense the cellular Pi status and binds to the intracellular Pi sensor SPX1 to control Pi homeostasis in plants [24]. We found that in addition to Pi homeostasis, the *TaVIH2-3B* promoter also responds to drought conditions.

Earlier, the double mutants of VIH genes in *Arabidopsis* show severe growth defects, implicating their unexplored role in overall growth and development[23]. We hypothesize that the molecular and biochemical changes in transgenic *Arabidopsis* provide the overall mechanical strength to the plant cell and, in turn, tolerance to stress conditions. These observations were also supported by our transcriptome analysis of two independent TaVIH2-3B overexpressing *Arabidopsis* lines that show consistent high expression of cell-wall, ABA and DREB genes (Figure 8 and Figure 9B). Multiple genes were differentially regulated by *TaVIH2-3B* overexpression, suggesting that increased protein levels of VIH2 could cause changes in gene expression patterns. Classically, VIH proteins contains evolutionarily conserved two distinct domains, including an N terminal rimK/ATP GRASP kinase and phosphatase domain. It remains to be dissected if the change in transcriptome response in these transgenic *Arabidopsis* is due to the kinase or phosphatase domain. Earlier, multiple inositol-1,3,4 tris*kis*phosphate 5/6-kinase (devoid of phosphatase domain) was also implicated for their role in drought tolerance[38, 39]. This may suggest that the tolerance for the drought could arise by the presence of the functional kinase domain.

Multiple studies have implicated that an enhanced level of ABA leads to drought tolerance[40–43]. The elevated levels of ABA in our transgenic plants could be accounted for the high expression of genes involved in cell-wall maintenance and biosynthesis. In yeast role of inositol pyrophosphate kinase was also implicated in vacuolar morphology and cell-wall integrity[14]. Plant cell-wall related remodelling and ABA regulated signalling is the primary response against abiotic stress, including drought[41, 42, 44]. ABA-dependent increased expression of NCEDs, *CYP*s and *DREBP* have been reported earlier in plants with their role implicated in drought stress[40, 43, 45]. Our study shows a high basal expression of genes encoding for DREBP and CYPs (Figure 8C). The high constitutive expression of these gene families in our transgenic *Arabidopsis* could account for their better adaptability for drought stress (Figure 8A, B and C). ABA is an important phytohormone regulating plant growth, development, and stress responses [46, 47]. At the mechanistic level ABA could target downstream genes that are able to support plant growth even under non-stress condition [48]. In our case high ABA levels could be as a result of such homeostatic interaction with other hormones; although this needs to be confirm in future. Additionally, the high expression of the sub-set of NCEDs and DREB genes could also be accounted for ABA regulated signalling in transgenic *Arabidopsis*. Similarly, overexpression of NCED could result in high accumulation of ABA[49, 50]. Earlier, changes in cellular levels of InsP_7_ and InsP_8_ have been attributed to guard cell signalling, ABA sensitivity and resistance to drought in maise *mrp5* mutants[31, 51]. This suggests a molecular link between TaVIH2, ABA levels and drought resistance. Resolving the in-vivo levels of InsPx is technically challenging for non-specialized labs. Our current study is limited due to the lack of in vivo measurements of InsP_8_ in these transgenic lines. In-vivo profiling of InsPx by enrichment with TiO_2_ is a powerful tool that has been employed with plant tissue [23, 24, 52]. We are currently optimizing this method to detect the InsPx generated in our transgenic lines. However, TaVIH2-3B showed the highest homology to AtVIH2 (70.6 %) and both show PP-IP5K like activity. Therefore, we speculate that these transgenic plants may possess high levels of InsP_8_.

*Atvih2-3* mutant lines lacking mRNA expression also show alteration in the cell-wall composition despite its typical growth as wild type Col-0 (Figure 9D). Interestingly, *vih1* and *vih2* double mutants display severe growth defect that was rescued by the gene complementation [23]. In our study, we complemented the *vih2-3* Arabidopsis mutant with the TaVIH2-3B that resulted in restoring Col-0(Ev) like phenotype. This suggests that wheat VIH2-3B could functionally complement Arabidopsis *vih2* mutants, and it is possible that the in-vivo level of InsP_8_ is restored in these lines since both bear PPIP5K activity.

Our overexpression data showing enhanced branching and robust growth collectively reinforce the notion that VIH are also involved in providing support for plant growth. The *vih2* mutant in *Arabidopsis* is more susceptible to infestation by caterpillar (*Pieris rapae*) and thrips[22]. The resistance against herbivore pathogens such as *P. rapae* could be gained by modulating the genes associated with cell-wall modification[53]. *Arabidopsis vih2* lines showed compositional changes in the cell-wall extracted polysaccharides, especially at the AG level. The decreased resistance in *vih2* mutants against herbivores could be accounted for the defect in the signalling pathway via COI1-dependent gene regulation and changes in the structural composition of the cell-wall. Taken together, we propose a working model, where wheat VIH participate in the drought resistance in plants by modulating the changes in cell-wall gene expression, enhanced ABA levels and change in biochemical composition to provide more mechanical strength (Figure 10E). In future, it will be interesting to quantitate the level of higher inositol pyrophosphates in these plants.

## Conclusions

Herein, we explored additional roles offered by plant VIH proteins. We employed genetic and biochemical tools to characterize the wheat homoeolog VIH2-3B as an active PP-IP5K. Our lines of evidence suggest that the expression of VIH genes is perturbed during drought conditions and could modulate the expression of genes involved in cell-wall maintenance so as to relay resistance to both mimic-drought and drought conditions. Interestingly, the wheat VIH2 was able to complement the *vih2-3/2-4* which were also sensitive to mimic-drought like condition. In summary, our work provides a glimpse into the emerging new role of plant VIH proteins in cell-wall scaffolding functions to provide resistance against drought stress. Future studies will be required to dissect the casual effect of drought response that could be mediated at the protein level by the VIH2 or levels of InsPx species in these transgenic lines.

## Methods

### Plant materials and growth conditions

The experiments in this study were conducted using *Arabidopsis thaliana* variety Col-0 ecotype and Bread wheat (*Triticum aestivum* L.) variety “C-306” (Mishra et al., 20201). For the collection of the tissue materials, the spikes tagged on the first day after anthesis (DAA), post which samples were collected at 7, 14, 21 and 28 DAA stages and various tissues, including root, stem, leaf and flag leaf of 14 DAA stage. For seed tissue collection 14 DAA seed was used to separate different tissues, including aleurone, endosperm, embryo, glumes and rachis as mentioned previously [54].

### Identification and cloning of two wheat VIH genes

Two *Arabidopsis* (AT5G15070.2 and AT3G01310.2) and the previously reported yeast VIP1 sequences were used to perform Blastx analysis against the IWGSC (www.wheatgenome.org/). The identified sequences were analyzed for the presence of the typical dual-domain structure. Furthermore, the Pfam domain identifiers of the signature ATP-Grasp Kinase (PF08443) and Histidine Acid Phosphatase (PF00328) domains were used to identify VIH proteins in the Ensembl database using the BioMart application. The corresponding predicted homoeologous transcripts were found and compared to the other VIH sequences. DNA STAR Lasergene 11 Core Suite was used to perform the multiple sequence alignment and calculate the sequence similarity. Gene-specific primers capable of amplifying the transcript from the specific genome was designed after performing 5’and 3’- RACE to ascertain the completed open reading frame (ORF). Subsequently, full-length primers were designed to amplify the *VIH* genes. The generated full-length sequence information was further used for qRT-PCR related studies.

### Hydropathy plot and IDR prediction

The hydropathy profile for proteins was calculated according to Kyte and Doolittle., 1982. The positive values indicate hydrophobic domains, and negative values represent hydrophilic regions of the amino acid residues. To identify the % similarity with IDR boundaries, MFDp2 (http://biomine.cs.vcu.edu/servers/MFDp2 was used to predict the disorder content in the input sequence[55].

### Isolation of total RNA, cDNA synthesis and quantitative real-time PCR analysis

Total RNA from various tissues was extracted by a manual method using TRIzol® Reagent (Invitrogen™). The integrity and concentration of RNA were measured, and contamination of genomic DNA was removed by subjecting the RNA samples to DNase treatment using TURBO™ DNase (Ambion, Life Technologies). 2 µg of total RNA was used for cDNA preparation using The Invitrogen SuperScript III First-Strand Synthesis System SuperMix (Thermo Fisher Scientific) as per the manufacturer’s guidelines. qRT-PCR was performed using the QuantiTect SYBR Green RT-PCR Kit (Qiagen, Germany). The gene-specific primers capable of amplifying 150-250 bp region from all the three homoeologous of two *TaVIH* genes were carefully designed using Oligocalc software. Four technical replicates for each set of primers and a minimum of two to three experimental replicates were used to validate the experiment. Gene-specific primer (with similar primer efficiencies) used in the study are listed in Supplementary Table S4. ADP-ribosylation factor gene (*TaARF*) was used as an internal control in all the expression studies. The Ct values obtained after the run were normalized against the internal control, and relative expression was quantified using the 2^−ΔΔ*C*^T method [56].

For In-silico expression for *TaVIH* genes, the RefSeq IDs were used to extract expression values as TPMs from the expVIP database. For different tissues and stages, the expression values were used to build a heatmap. In the case of abiotic and biotic stress conditions, the expression values from the control and stressed conditions were used to get fold change values, which were then used to plot heatmaps using MeV software.

### Construct preparation for expression vector and yeast functional complementation

For complementation assays, pYES2, a galactose-inducible yeast expression vector, was used. The functional complementation of yeast by TaVIH proteins (with C-myc tag) was studied using 6-azauracil based assay. The wild type BY4741 (MATa; his3D1; leu2D0; met15D0; ura3D0) and *vip1*Δ (BY4741; MATa; ura3Δ0; leu2Δ0; his3Δ1; met15Δ0; YLR410w::kanMX4) yeast strains were used for the growth assays. The CDS corresponding to the catalytic domain of *ScVIP1* (1-535 amino acids) cloned into pYES2 expression vector was used as a positive control. *TaVIH1/2*, along with *ScVIP1* and empty vector, were transformed individually into wild type and mutant strains by the lithium acetate method with slight modifications. The expression of both TaVIH1-4D and TaVIH2-3B in yeast was confirmed by Western blotting using Anti C-myc antibody (1:1000; raised in mice; Invitrogen, USA). For growth assay, the wild type and mutant *S. cerevisiae* strains carrying different plasmids were allowed to grow overnight in minimal media without uracil. The primary culture was used to re-inoculate fresh media to an OD_600_ of 0.1 and grow until the culture attained an optical density of 0.6-0.8. The cell cultures were then adjusted to O.D of 1 and further serially diluted to the final concentrations of 1:10, 1:100 and 1:1000. 10 μl each of these cell suspensions were used for spotting on SD(-Ura) plates containing 2% galactose, 1% raffinose and varying concentrations of 6-azauracil (0, 2.5 and 5 mM). The colony plates were incubated at 30°C, and pictures were taken after four days.

### Protein expression of wheat VIH1 and VIH2, In-Vitro Kinase assays, PAGE analysis and MADLI-ToF analysis

The TaVIH1-KD and TaVIH2-KD were cloned in pET-28a and expressed in *E. coli* BL21 cells using 0.5 mM IPTG and purified in lysis buffer having pH 7.4 containing 50 mM sodium phosphate, 300 mM NaCl and protein inhibitor cocktail. Post sonication and centrifugation purification was done on the Cobalt resin affinity chromatography column (ThermoFisher Scientific, Waltham, MA, USA). After column saturation overnight at 4°C, it was washed with buffer containing 7.5 mM imidazole and subsequently eluted with buffer containing 100 mM EDTA. The eluate was pooled and concentrated using a concentrator having a molecular weight cut-off of 10 kDa by spinning at conditions mentioned in the vivaspin concentrator’s manual. The concentrated enzyme preparation was washed thrice with sodium phosphate buffer and finally concentrated in Tris-HCl buffer, pH 7.4. Purified proteins were analyzed by western blotting with Mouse anti-HIS primary antibody and Goat anti-Mouse secondary antibody [HRP IgG (H + L): 1:5000 dilutions; Invitrogen].

Kinase assays were performed using the ADP-Glo ™ Max Assay kit (Promega, USA) according to the manufacturer’s guidelines. This kit utilizes the luminescence-based test for ADP quantification as a measure of kinase activity. We prepared InsP_7_ by using 100ng of Mouse IP6K1 (mIP6K1) recombinant protein along with 100 µM of InsP_6_ (Sigma, USA) in a buffer containing 20 mM HEPES (pH 7.5), 5 mM MgCl_2_, 10 mM ATP and 1 mM DTT for 3 hr at 28°C. The resultant product was first resolved by TBE-PAGE gel and then eluted from the gel as described earlier [57] and was used for the reaction. The concentration of the eluted InsP_7_ was measured with ImageJ software by comparing with varying InsP_6_ concentrations in the TBE PAGE gels[57]. For ADP-Glo ™ Max Assay kit 50 ng of respective protein (VIH1 and VIH2) and 300 nM InsP_7_ and 1µM of ATP was used, and the assay was conducted by following the manufacturer’s guidelines. Luminescence was measured one hour after adding the ADP-Glo™ Max Detection Reagent, using SpectraMax M5e plate reader (Molecular Devices, USA).

For resolving the InsPx species generated by TaVIH1 and TaVIH2, separate kinase assays were performed in 20 mM HEPES (pH 7.5), 5 mM MgCl_2_, 10 mM ATP, 100 µM InsP_7_ and 1 mM DTT and incubated along with 30 ng of respective proteins in a total volume of 100µl. ScVIP1 was taken as a control for the reaction. These reactions were incubated at 28°C for 1, 2 or 9 hrs. The reaction products were separated by TBE-PAGE and visualized by Toluidine Blue staining. All the inositol polyphosphates were resolved using 18 cm gel using 33.3 % polyacrylamide gel in Tris-Borate EDTA, as mentioned earlier [28]. These gels were pre-run for 75 min at 300 volts, and the samples were mixed with dye (10 mM Tris-HCl pH 7.0; 1 mM EDTA; 30 % glycerol; 0.08 % Orange G) and loaded. Gels were run at 5-6 mA overnight at 4°C until the Orange G dye front reached 6 cm from the bottom of the gel. Bands were subsequently visualized by Toluidine Blue (0.1 % w/v) stain. TBE-PAGE gel-purified products of TaVIH reaction were used for Matrix-assisted laser desorption-Time Of flight Mass Spectrometry analysis (MALDI-ToF-MS). MALDI-ToF-MS was performed from gel extract solutions which were pipetted on an α-Cyano-4-hydroxycinnamic acid (≥98%, Sigma) prepared on a stainless-steel plate (0.5 μL of a 10 mg/mL ACN/H2O 1:1 solution). Negative ionization mode was used for acquiring spectra on a spectrometer (AB SCIEX TOF/TOF^TM^ 5800; equipped with a 337 nm laser) operating in the linear mode.

### Cloning of VIH promoter, cDNA and *Arabidopsis* transformation

For promoter, ∼2000 bp fragments upstream of the start codon were PCR amplified from genomic DNA. The cloned DNA fragments (in pJET1.2) were sequenced, confirmed and inserted into pCAMBIA1391z, a promoter-less binary vector containing GUS reporter gene to generate *TaVIHpromoter: GUS* in pCAMBIA1391z. For *VIH2-3B* cDNA (3117 bp) fussed C-terminal His tag, site-directed cloning was done at *Spe*1 generated site in pCAMBIA1302 (pCAMBIA1302:*TaVIH*-His). These generated transcription units were introduced into *Arabidopsis* seedlings, or T-DNA insertion lines of *vih2-3* (SAIL_165_F12), *vih2-4* (GK-080A07) mutant using *Agrobacterium tumefaciens* (GV3101) mediated transformation by floral dip method (Zhang, Henriques, Lin, Niu & Chua 2006). Multiple (7-10) independent transformants were screened on 0.5X MS media containing 30 mg/L hygromycin and 0.8% agar. The transformed seedlings with long hypocotyls and green expanded leaves at a 4-leaf stage were separated from the non-transformed seedlings and transferred to the soil after about three weeks. Similarly, T_1_ and T_2_ generation seeds were also selected and allowed to grow till maturity. The transgenic seedlings were confirmed for the presence of recombinant cassette using PCR based approach. The transgenic lines harbouring empty pCAMBIA1391Z or pCAMBIA1302 vector was used as a respective negative control. The PCR positive lines were further used for functional characterization. In addition, the promoter sequences of *TaVIH* genes were analyzed for the presence of cis-regulatory elements using the PLANTCARE database (http://bioinformatics.psb.ugent.be/webtools/plantcare/).

### GUS-reporter assays and characterization of transgenic lines *in Arabidopsis*

For promoter analysis, the seeds of PCR positive lines were surface sterilized and grown on 0.5X MS (Murashige and Skoog media) agar plates containing 30 mg/L Hygromycin B for 15 days before they were subjected to various abiotic stress and hormonal treatments. For dehydration stress, the seedlings were air-dried by placing them on Whatman filter paper for 1hr. Exposure to ABA (100 µM), GA_3_ (20 µM) and drought-mimic (20% and 30% PEG) were given by placing the seedlings on filter paper impregnated with 0.5X MS solution containing the respective chemical for 24 hrs. For Pi starvation, seedlings were allowed to grow on MS agar plates without KH_2_PO_4_ for 96 hrs. Histochemical staining of seedlings after respective treatments were performed by incubated overnight in GUS staining solution (Jefferson 1987) with 2 mM X-Gluc (5-bromo-4-chloro-3-indolyl-beta-D-glucuronic acid, HiMedia, India) at 37°C in a 48-well microplate containing about ten seedlings/well. Chlorophyll was removed from tissues by dipping in 90% ethanol. The staining was visualized and photographed under Leica DFC295 stereomicroscope (Wetzlar, Germany) at a magnification of 63X. MS solution without any chemical served as a control.

For characterization of transgenic lines parameters such as rosette area, the number of leaves, leaves size, length of central root axis and number of shoots (primary and secondary). Four independent confirmed homozygous transgenic lines were used for this study. Each parameter was calculated using three experimental replicates, each consisting of twelve plants. For drought-mimic stress experiments, three days old seedlings of transgenic and control pre-grown on 0.5X MS were transferred to 0.5X MS plates consisting of either 125-or 100-mM mannitol or 5 or 10 % glycerol. Ten seedlings were used, and the experiments were repeated three times with similar phenotypes. For control, seedlings continued to grow on ½ MS plates. Root lengths were measured, graphs were plotted (using three experimental replicates), and pictures were taken after nine days of growth. The relative water loss % was calculated of twenty-five leaves per five plants with a similar developmental stage for each of the transgenic lines, and control plants were subjected to incubation (27°C) for the period of 8 hrs. The fresh weight of the detached leaf was taken and continued for the measurements every 2 hrs. The experiment was repeated twice for similar observations. The leaf relative water content (RWC) measurement was performed as mentioned earlier [41]. The value for each treatment was calculated by using the standard formula RWC (%) = [(FW-DW)/(TW-DW)] X 100 with FW is fresh weight, DW is dry weight, and TW is turgor weight. For performing these experiments, leaves of equal sizes were detached from 24 days old transgenic lines and control *Arabidopsis* and weighed immediately (FW). The leaves were submerged in deionized water for 24 hrs. After incubation, the leaves were blotted dry, and their weight was determined (TW). To measure their DW, they were oven-dried (at 65°C) for 24 hrs. The experiments were performed with at least three experimental replicates, each consisting of five to six plants.

For drought response, the seedlings were grown in a symmetrical box with demarcated sections for each seedling. The seedlings were inter-rooted so that they are exposed to similar soil moisture conditions. The seven days old seedlings of *Arabidopsis* were subjected to drought (water-withholding) conditions for the period of fourteen days. After the drought period, the seedlings were re-watered, and observations were made after 4 and 7 days. Post this, the plants were observed, and % survival rates were calculated.

### RNAseq profiling

Col-0(Ev) and overexpressing *TaVIH2-3B* Arabidopsis (#Line4 and 6) seedlings were grown for 25 days. Total RNA was extracted from three independent biological replicates for each genotype using RNeasy Plant Mini Kits (Qiagen, CA). Genomic DNA contamination was removed by digestion with Turbo DNase (BioRad, CA). RNA quantity was checked by Bioanalyzer for quality control (RIN>8). Library construction and sequencing were performed by Eurofins, Bangalore, India, using pair-end library preparation. About 9.5 to 13.8 million raw reads were obtained for each sample. Raw reads were processed to filter out the adapter, and low quality (QV<20) reads using trimmomatic v0.39[58]. The reads were then pseudo-aligned against the reference transcriptome (Ensembl release 48) using Kallisto v0.46.2 [59]. The obtained raw abundances were summarised to gene-level expression counts using tximport and imported to DESeq2 [60, 61] for differential expression (DE) analysis in R. The obtained log2 fold change (LFC) values were further processed using apeglm package to reduce noise [26]. Genes with 1 > LFC < −1 and padj < 0.05 were considered significantly DE. The expression correlation across lines and within replicates was analyzed using Principal Component Analysis (PCA) in ggplot2 [62]. The data have been deposited in the NCBI as a Bioproject ID PRJNA685929.

### GC-MS analysis of Arabidopsis cell-wall polysaccharides and ABA measurement

Extraction of cell-wall components was performed as described earlier with minor modification as depicted in the flowchart as Supplementary Figure S10 [63]. Since such chemical analysis requires relatively large amounts of samples, pools from 3-5 independent plants (for each of the three biological replicates) of the respective lines expressing wheat VIH2-3B were used for chemical analysis. Briefly, five grams (fresh weight) of shoots from respective lines and control at similar developmental stages (25 days old) was crushed to a fine powder and processed further. The derived pellet was used to extract arabinoxylan (AX) and cellulose, whereas the supernatant was used to extract arabinogalactan (AG). The extractions were checked with Iodine solution to make sure that they are free of starch interference. The compositional analysis of the extracted AG, AX and Cellulose was determined by preparing their alditol derivatives and process for gas chromatography-mass spectrometry (GC-MS) analysis as described [64, 65]. Two µl of samples were introduced in the splitless injection mode in DB-5 (60 m × 0.25 mm, 1 µm film thickness, Agilent, CA) using helium as a carrier gas. The alditol acetate derivative was separated using the following temperature gradient: 80 °C for 2 min, 80-170 °C at 30°C/min, 170-240 °C at 4°C/min, 240°C held for 30 min and the samples were ionized by electrons impact at 70 eV. ABA was measured using Plant Hormone Abscisic Acid (ABA) ELISA kit (Real Gene, Germany). Twenty-five days old plants leaves were used for the measurement of the ABA content. One gram of fresh weight from eight plants for each line was used for extractions. The experiments were repeated with at least three independent extractions, and concentration was calculated using standard graphs as per the manual instructions. Standard graph and test samples were plotted using a log of concentration, and colour development for each line was measured at 430 nm (Supplementary Figure S11A and B).

## Acknowledgements

Authors thank Executive Director for facilities and support. Part of this work was also supported by the NABI-CORE grant to AKP. MK thanks UGC-CSIR for her research scholarship. Thanks to Dr Gabriel Schaff for sharing the *Arabidopsis vih2-3* and *vih2-4* T-DNA insertion mutant. AS thank DBT for the SRF fellowship. DBT-eLibrary Consortium (DeLCON) is acknowledged for providing timely support and access to e-resources for this work.

## Availability of data and materials

All data generated or analyzed during this study are included in this published article and its supplementary information files. The resources, including plasmids, constructs, and transgenic *Arabidopsis* seeds, will be available upon reasonable request.

## Contributions

AS, AKP, RB, PP, MB, HR and VR wrote the text and drafted the figures; AS, SK, KM, AKP, MB, RB, SG and VK designed and conducted the establishment experiments; GK, AKP and AS conducted the RNAseq experiments and analyzed the data. AS, MB and SS performed reporter assays and transgenic related work; AS performed PAGE assays, RB and SG assisted in the analysis of inositol polyphosphates by PAGE. All authors read and approved the final manuscript. AKP acquired the funding from the extramural agency.

## Ethics approval and consent to participate

Not applicable.

## Consent for publication

Not applicable.

## Supplementary Figure

**Supplementary Figure S1:** Kyte-Doolittle Hydropathy plots and conserved domains of wheat VIH proteins. (A) Kyte-Doolittle hydropathy plots with the positive values indicating the hydrophobic domains and negative values represent hydrophilic regions of the amino acid residues. The hydropathy profile for proteins was calculated according to Kyte and Doolittle., 1982. (B) Schematic representation of domain architecture of TaVIHs deduced from CDD database: light grey rectangles indicate ATP Grasp/RimK Kinase domain and dark grey coloured hexagon corresponds to Histidine Phosphatase superfamily.

**Supplementary Figure S2:** Multiple Sequence Alignment (MSA) of different VIH/Vip protein sequences (TaVIH1, TaVIH2, AtVIH1, AtVIH2 and ScVIp1). The red sequence shows high conservation of the amino acids. The single green line indicates rimK/ATP-Grasp Kinase domain, and the double green line indicates Histidine Phosphatase Domains (HAPs).

**Supplementary Figure S3:** Yeast complementation assays of wheat VIHs. (A) Total protein was extracted from yeast cell transformed with TaVIH1-4D (C-myc tag) and TaVIH2-3B (C-myc tag) and Western analysis was done (left panel). Representative image of spotting assay performed on SD-Ura plates containing 1% raffinose, 2% galactose and supplemented with 0, 2.5 and 5mM of 6-azauracil (right panel). The wild type BY4741 and *vip1*Δ strains were transformed with respective constructs using Li-acetate method. Representative images were taken 4 days after the spotting assay was performed. Similar results were obtained with three independent repeats. (B) Filamentous growth assays were observed for wild type yeast (WT), yeast mutant-*vip1*Δ with empty pYES2 (*vip1*Δ) and *TaVIH2-3B* complementation in vip1Δ-(*TaVIH2-3B*+ Δvip1). Pictures were taken 20 days post-incubation.

**Supplementary Figure S4:** Protein purification and western analysis of wheat TaVIH1-KD and TaVIH2-KD. The molecular weight is around 40 kDa. Both the VIH proteins (VIH1 and VIH2) were expressed and purified as mentioned in the Methods section, and the expression was confirmed by the Western analysis using His-antibody.

**Supplementary Figure S5:** (A) PAGE-gel (33%) analysis of mIP6K1 generated product by staining with Toluidine Blue. Substrate InsP_6_ without and with ATP (2.5, 5 and 10 mM) was used as a control. The product InsP_7_ was generated using mIP6K1 and InsP_6_ as a substrate (last lane). (B) The InsP_7_ generated by mIP6K1 was eluted from gel and MS analysis was done which indicated a signal at m/z of 740.3 that correspond to mass of InsP_7_ and matches with the expected generated species. Indicated by arrow. (C) The kinase reactions were performed using 30 ng of TaVIH1-KD, and TaVIH2-KD purified proteins for 9 hr at 28° C. (D) MALDI-ToF MS analysis of synthesized InsP_8_ for TaVIH2-3B KD. MS analysis indicated a significant signal at m/z of 820.47 that correspond to the mass of InsP_8._ Indicated by arrow.

**Supplementary Figure S6:** Hormonal and abiotic stress response of *TaVIH* genes promoter. (A) Cis-element analysis of VIH1 and VIH2 promoters (∼1kb) Multiple stress related domains are represented in a schematic form. (B) Representative images for histochemical GUS assay performed against different stresses for promTaVIH1:GUS and promTaVIH2:GUS transgenic lines raised in *Arabidopsis thaliana* Col-0 background. Two week old seedlings selected positive against hygromycin selection on 0.5XMS agar plates were subjected to respective treatments: drought (20% PEG), dehydration (1hr air drying), ABA (100µM), GA_3_ (20µM) and Pi-deficiency (0.5X MS medias without KH_2_PO_4_). Seedlings with or without treatment (control) were stained overnight in GUS staining solution and photographed using Leica stereomicroscope at 6.3X magnification.

**Supplementary Figure S7: RNAseq analysis of transgenic Arabidopsis.** (A) PCA analysis of the RNAseq for control (Col-0 (Ev)) and two transgenic Arabidopsis lines. (B) Map man analysis of the genes those are consistently represented in the two transgenic Arabidopsis lines with overexpressing TaVIH2-3B.

**Supplementary Fig. S8:** qRT-PCR validation of selected genes from the Col-0(Ev), #Line4 and #Line6. A total of 2 µg of RNA (DNA free) was used for cDNA synthesis and qRT-PCR was performed using gene specific primers (Supplementary Table S4). C_t_ values were normalized against wheat *ARF1* as an internal control.

**Supplementary Figure S9:** Expression patterns of *TaVIH* gene homoeologous in different tissues and stress conditions. RNAseq datasets of (A) Tissues and developmental stages (B) Abiotic (phosphate starvation, heat and drought stress) and (C) Biotic stress conditions were used. The expression values were obtained from expVIP database in the form of TPM values and ratios of stressed to control condition were used to generate heatmaps using MeV software. Green and red colors represent down-regulation and up-regulation of the genes in the specific stresses, as shown by the color bar.

**Supplementary Figure S10:** Flow representation of the preparation and extraction of polysaccharides (Arabinogalactans, Arabinoxylans and Cellulose) form the shoots of *Arabidopsis*.

**Supplementary Figure S11:** Standard graph for ABA measurement in plant leaves samples. (A) Y-axis indicates Log of concentration and X-axis indicates the optical density. Data was linearised by plotting the log of the target antigen concentrations versus the log of the OD and the best fit line was determined by regression analysis. (B) Panel showing the colour development for the quantitation of the ABA in different leaf samples, OD was taken at 420 nm.

**Supplementary Table S1:** List of *TaVIH* genes with computed physical and chemical parameters. The molecular weight and isoelectric point prediction were done using Expasy ProtParam tool (https://web.expasy.org/protparam/). The sub-cellular localization prediction was done using WoLF PSORT prediction tool (http://www.genscript.com/wolf-psort.html). RefSeq v1.1 for wheat Ensembl Plants was used for gene ID.

**Supplementary Table S2:** List of genes up- and down-regulated in #line4 (Sheets1,2) and line6 (Sheets3,4) w.r.t. Col-0(Ev) lines. DEGs were obtained using the Kallisto-DESeq2 pipeline; genes with LFC > 1 in either direction and padj < 0.05 were considered to be differentially regulated.

**Supplementary Table S3:** List of drought responsive genes that are differentially regulated in #line4 (Sheet1), #line6 (Sheet2), and differentially regulated in both #line4 and line6 (Sheet3). Drought responsive genes at 10days of drought stress w.r.t Control plants were extracted from the SRA RNAseq dataset (SRP075287) using Cufflinks pipeline. Genes with 1 > LFC < −1 were considered to be drought responsive.

**Supplementary Table S4:** List of primers used for this study

