## Supplementary Figure Files for "Wheat inositol pyrophosphate kinase TaVIH2-3B modulates cell-wall composition for drought tolerance in Arabidopsis"

**Supplementary Figure S1:**

(A)

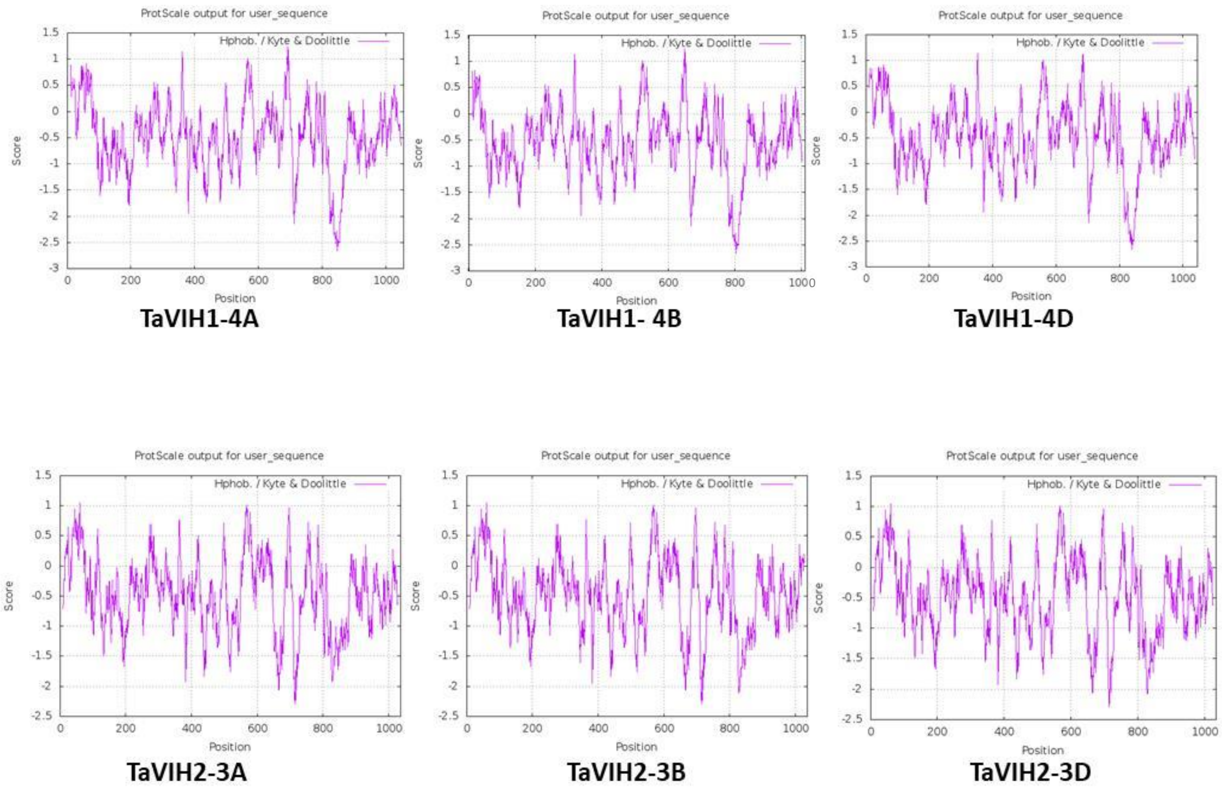


(B)

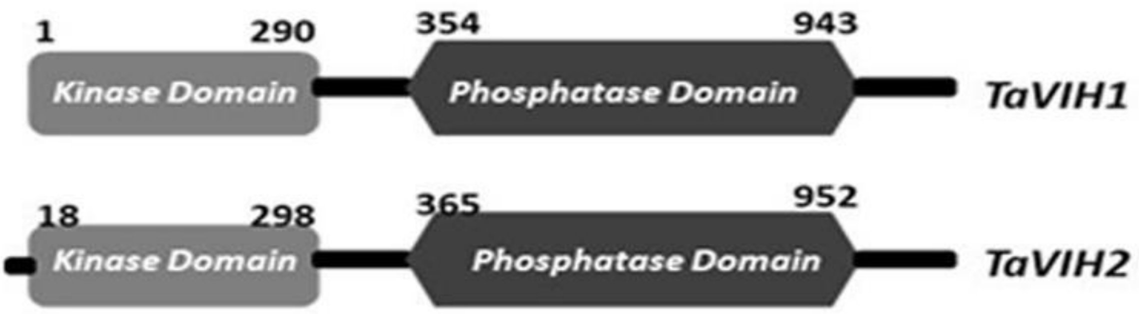

**Supplementary Figure S2:**

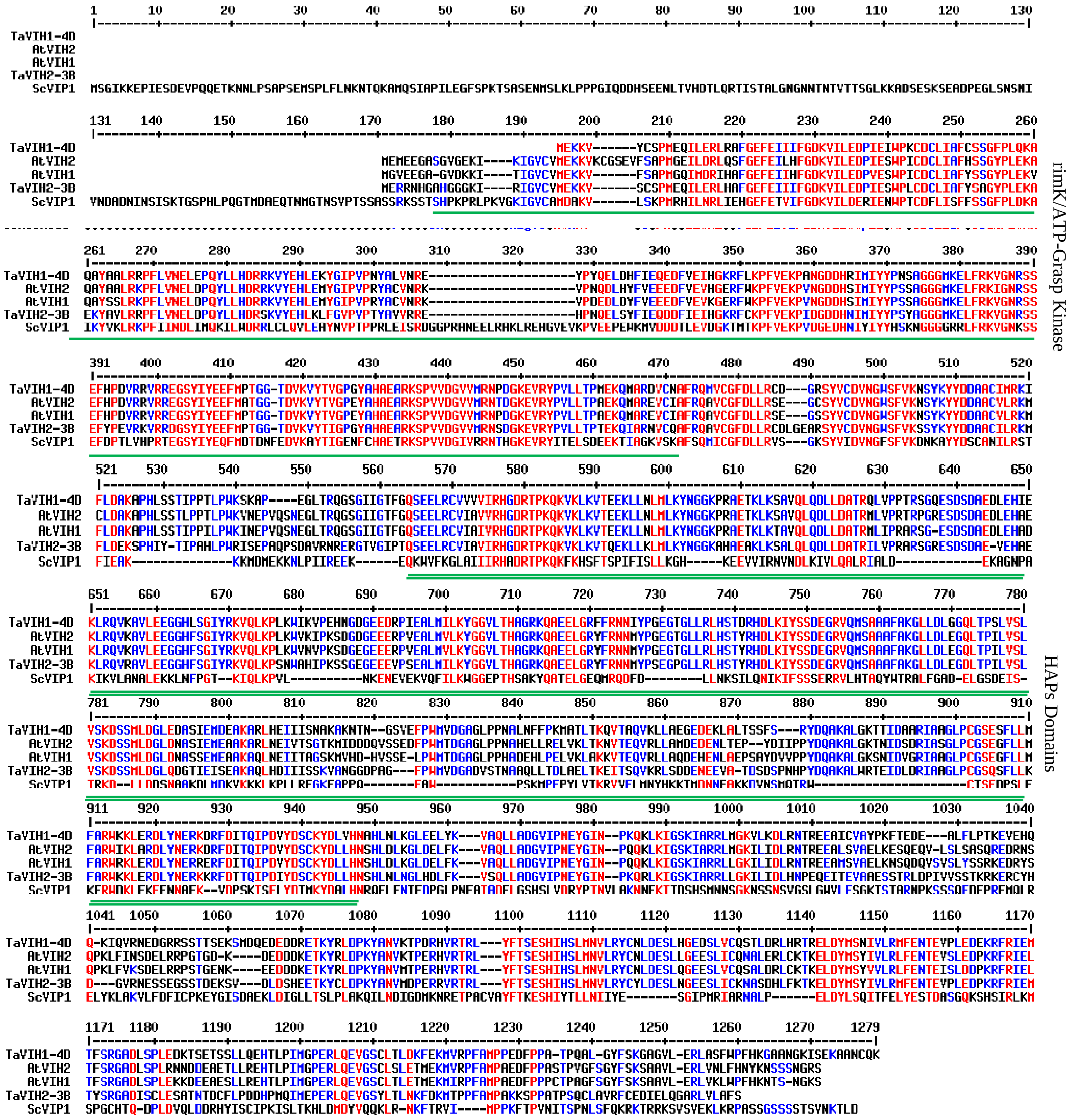

**Supplementary Figure S3:**

(A)

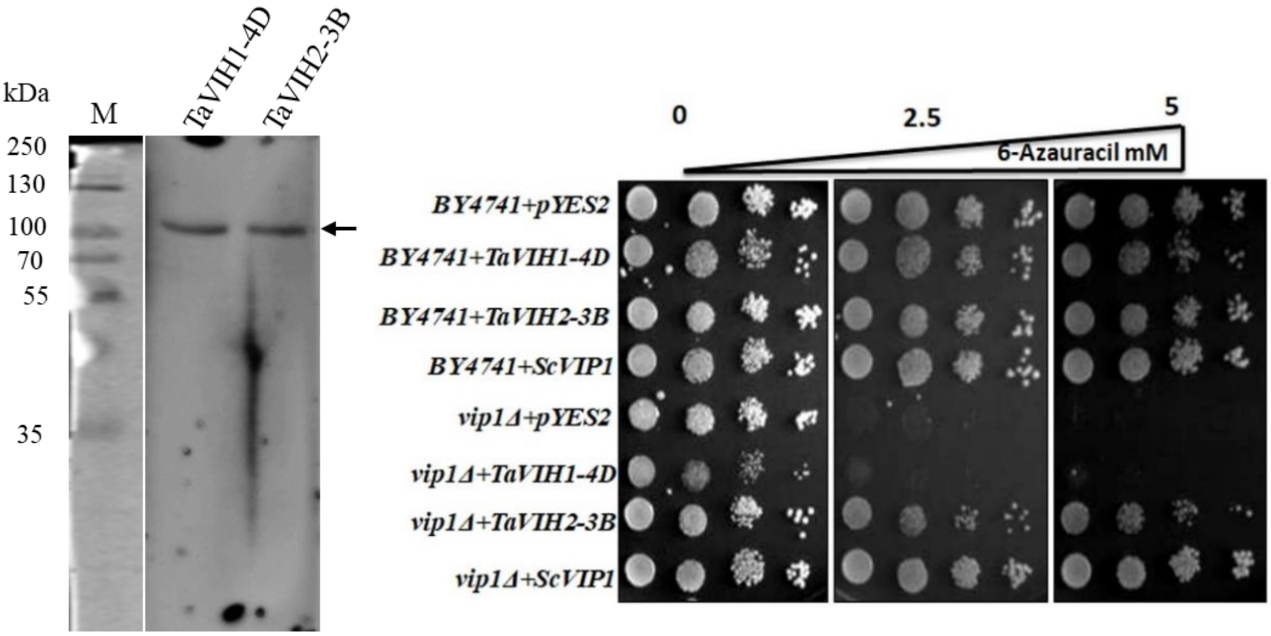


(B)

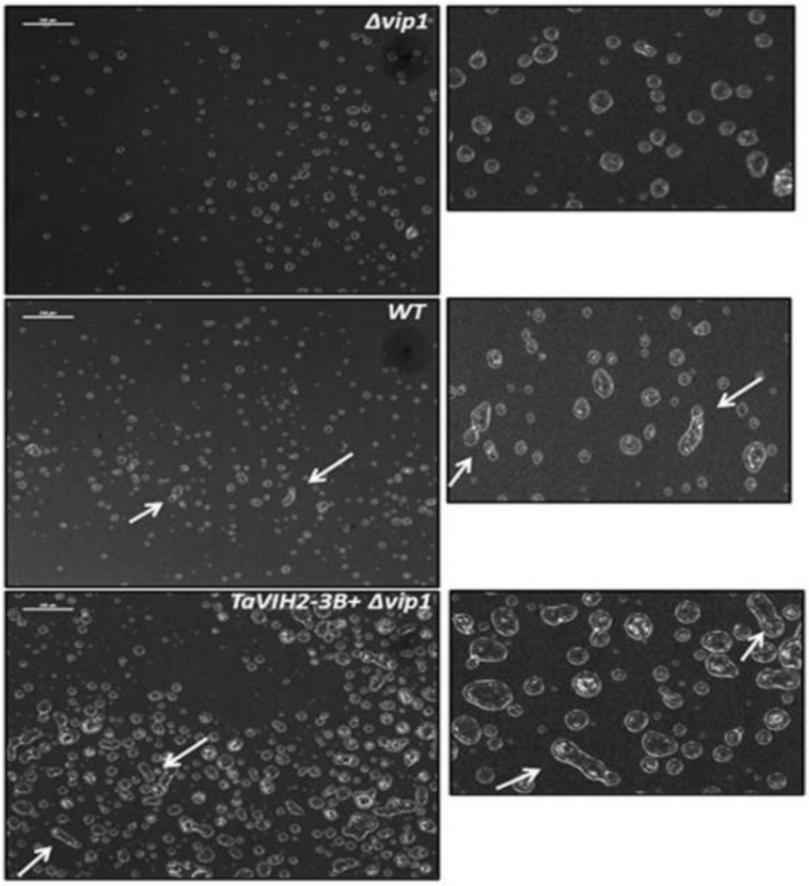

**Supplementary Figure S4:**

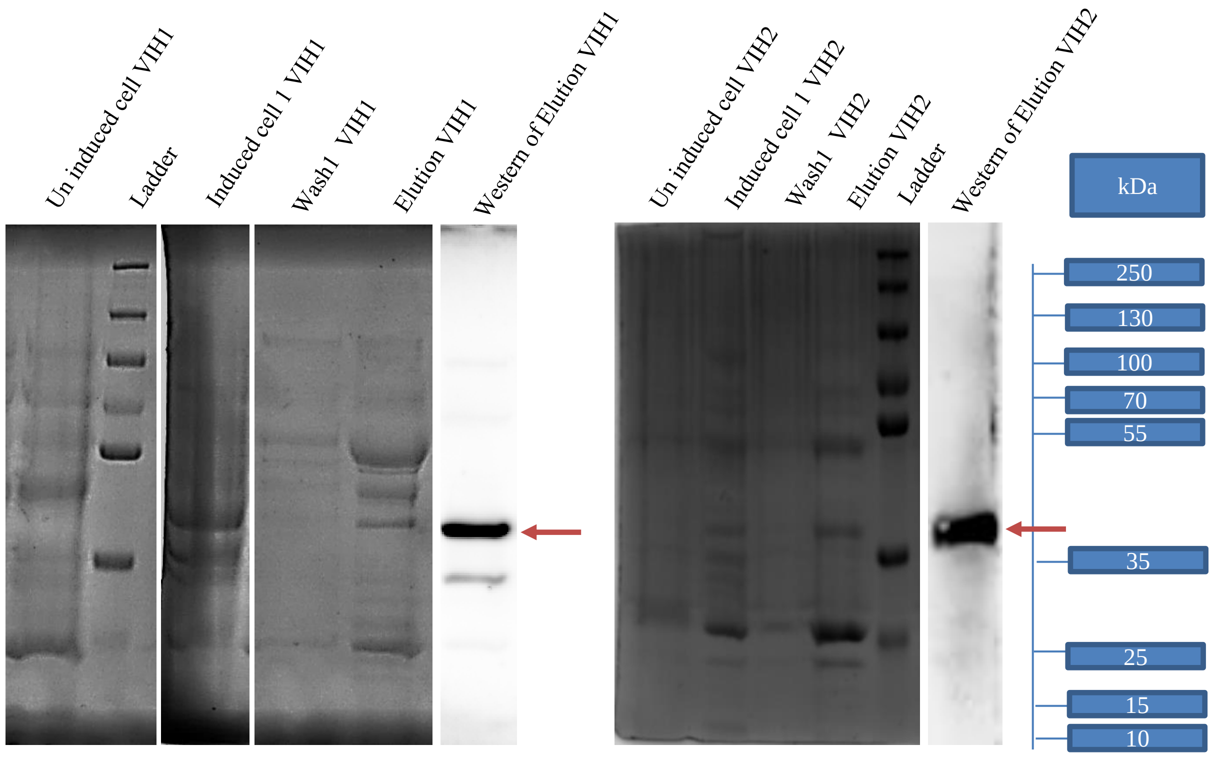

**Supplementary Figure S5:**

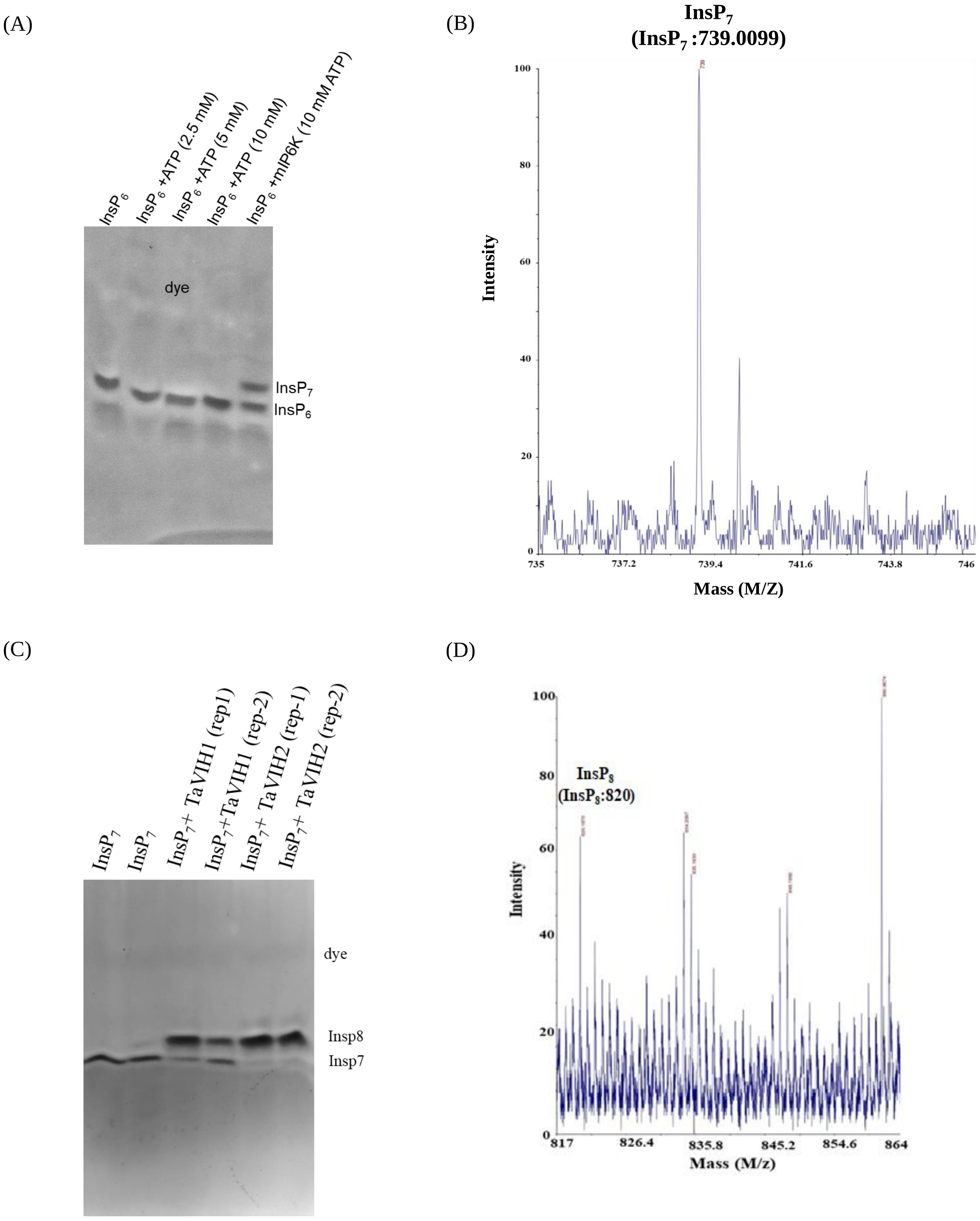

**Supplementary Figure S6:**

(A)

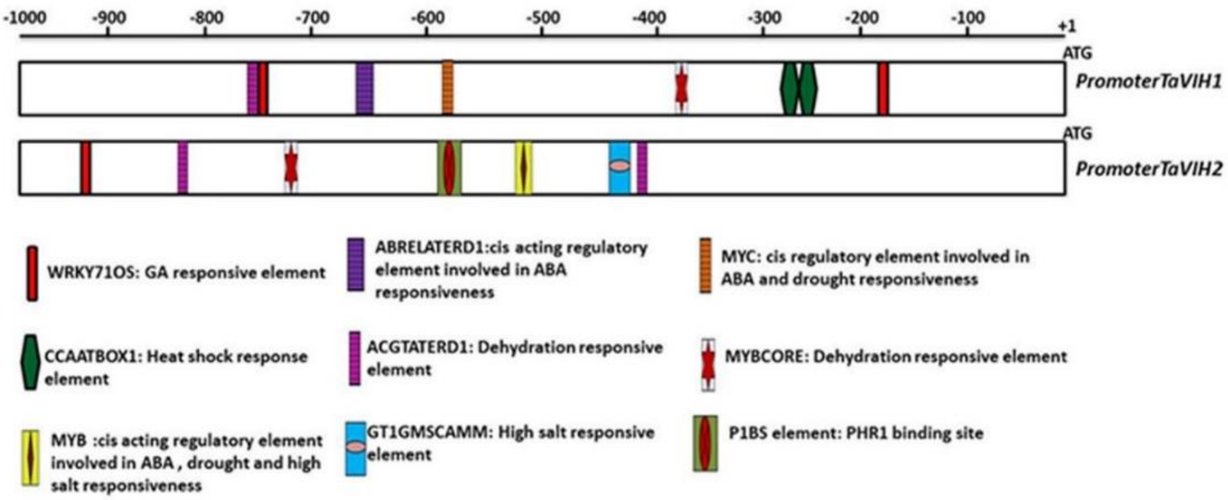


(B)

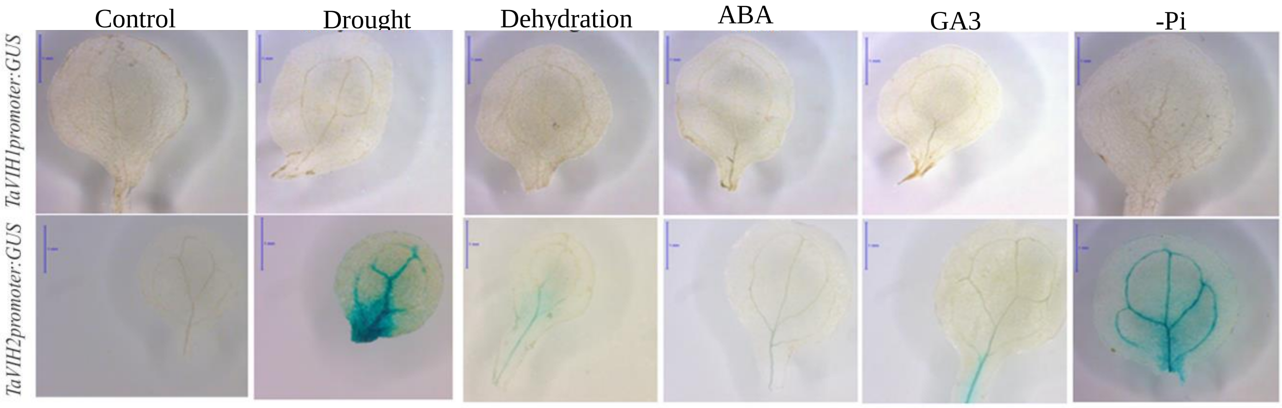

**Supplementary Figure S7:**

(A)

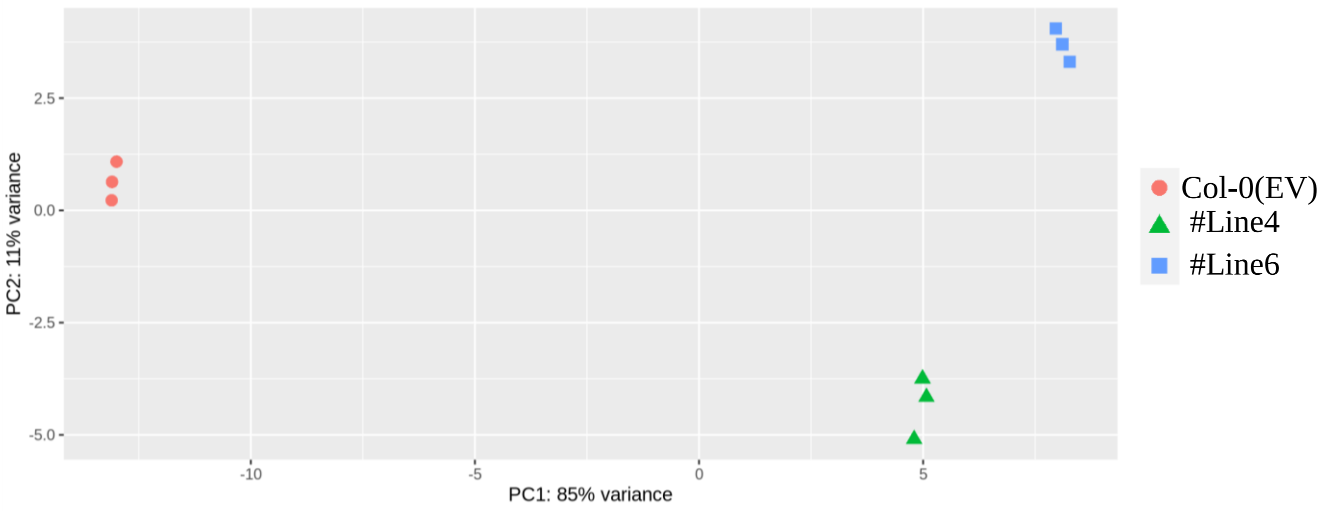


(B)

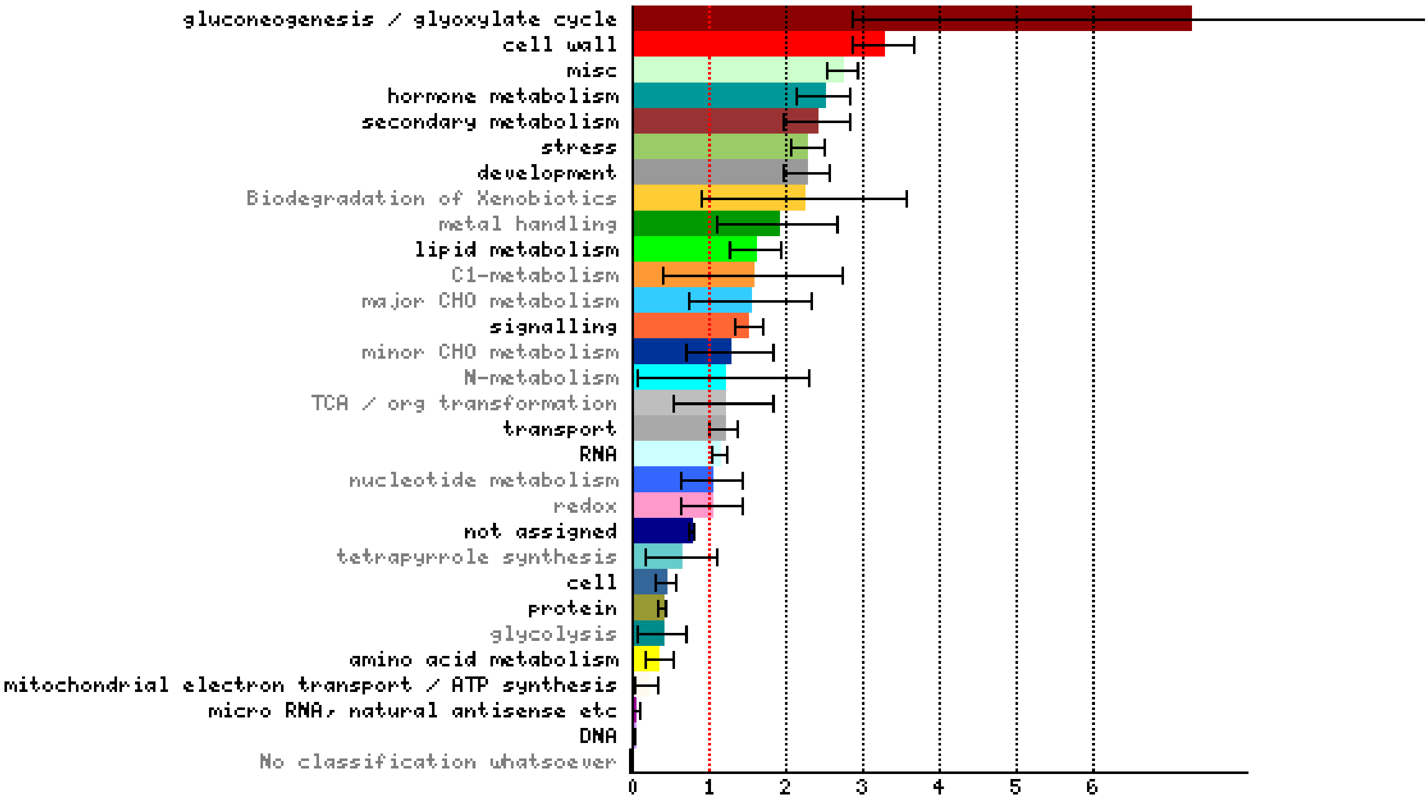

**Supplementary Figure S8:**

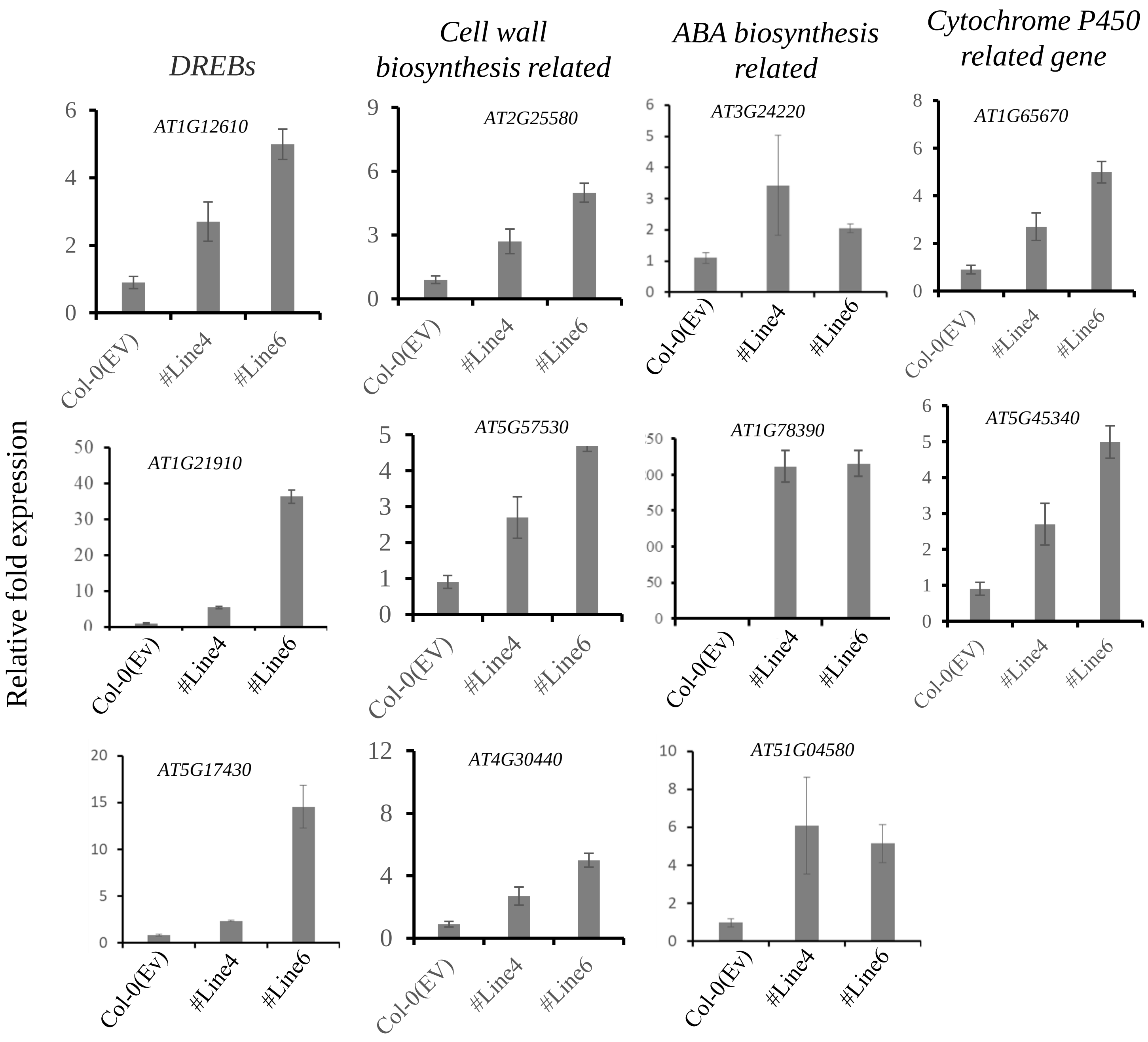

**Supplementary Figure S9:**

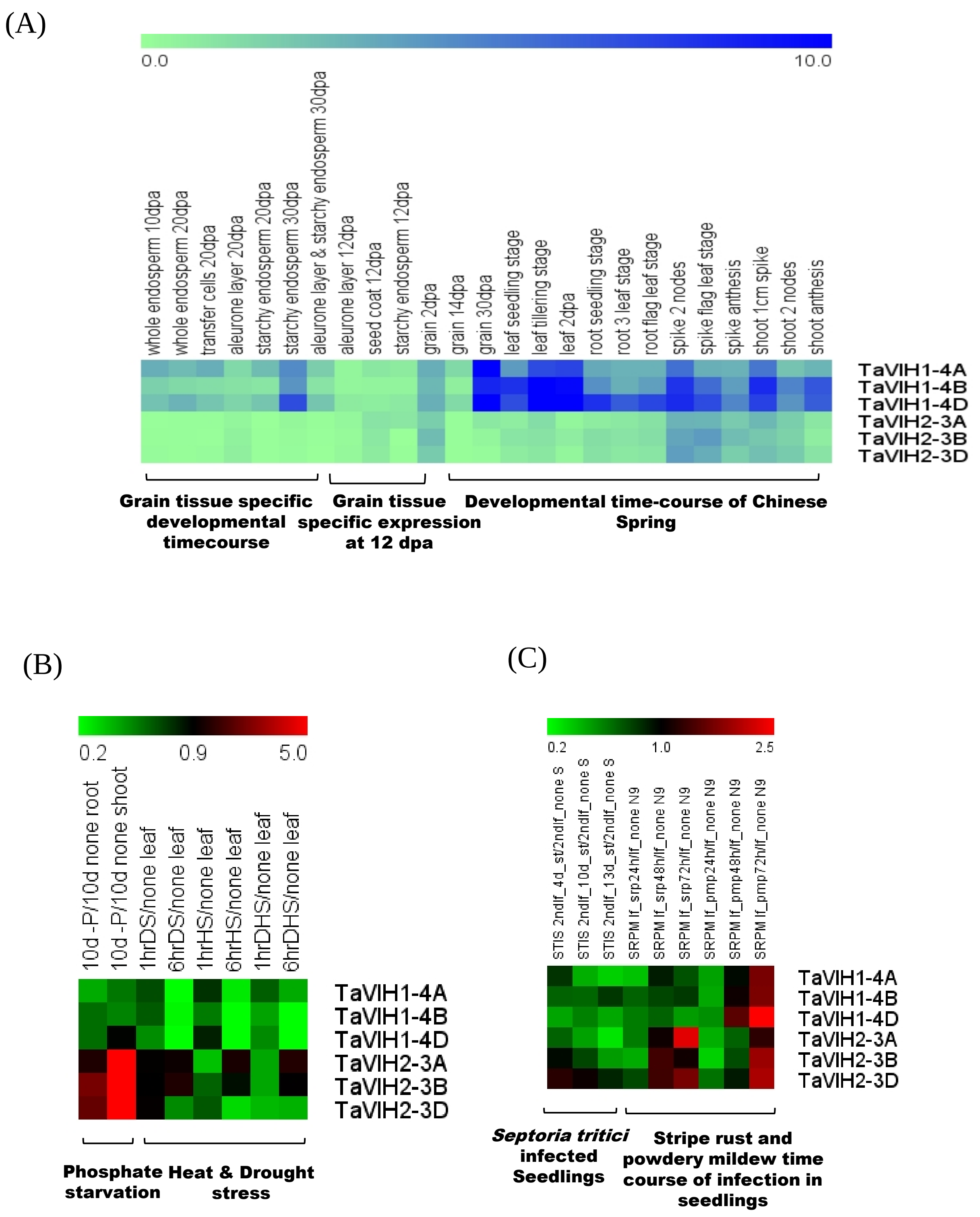

**Supplementary Figure S10:**

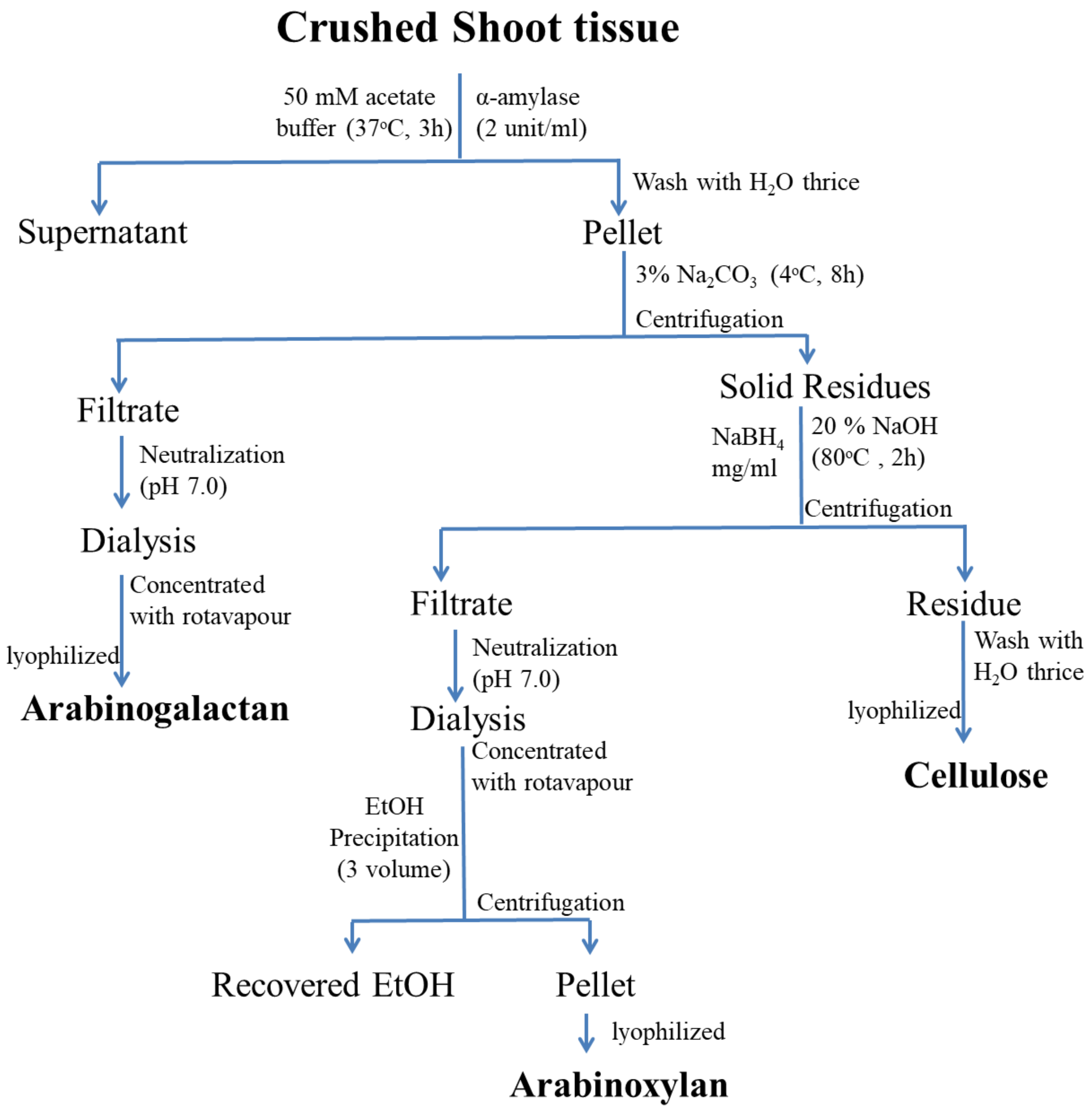

**Supplementary Figure S11:**

(A)

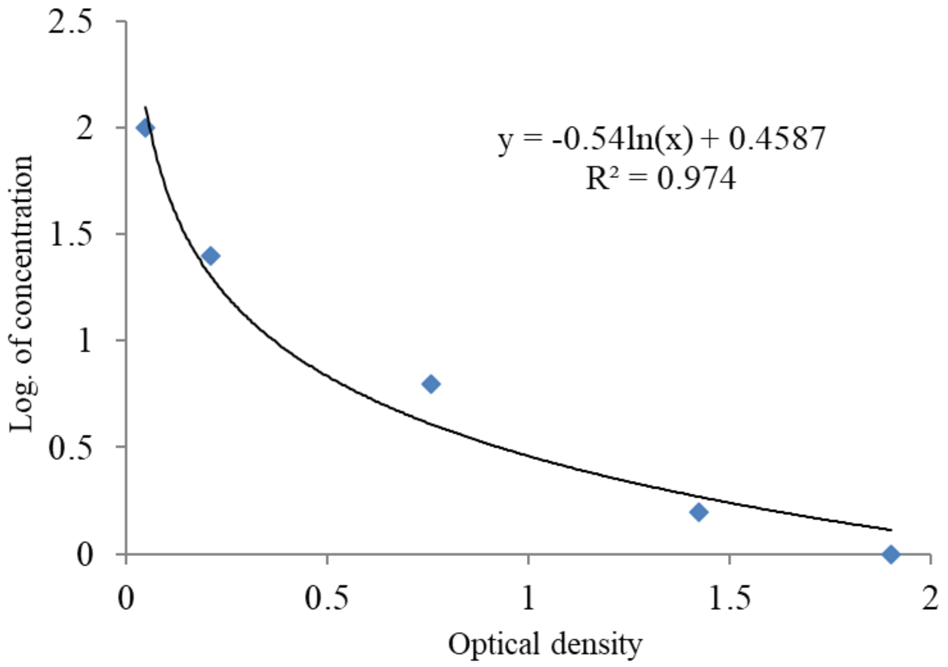


(B)

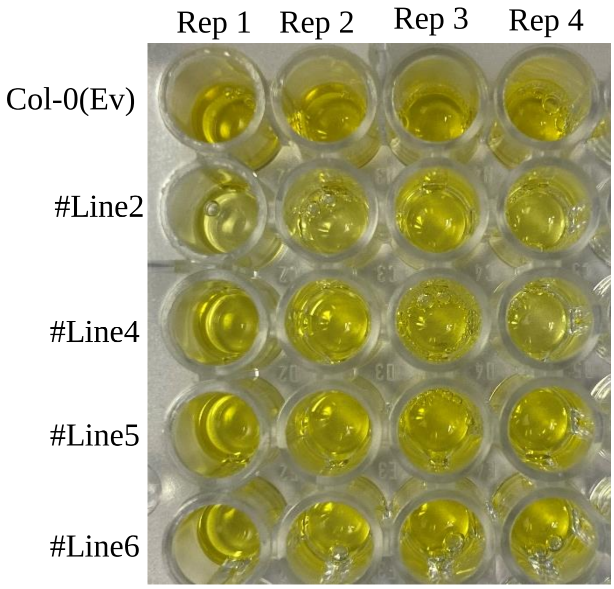
