## Supplementary material for "Wheat inositol pyrophosphate kinase TaVIH2-3B modulates cell-wall composition for drought tolerance in Arabidopsis": Table S1

**Supplementary Table S1:** List of *TaVIH* genes with computed physical and chemical parameters. The molecular weight and isoelectric point prediction were done using Expasy ProtParam tool (https://web.expasy.org/protparam/). The sub-cellular localization prediction was done using WoLF PSORT prediction tool (<http://www.genscript.com/wolf-psort.html>). RefSeq v1.1 for wheat Ensembl Plants was used for gene ID.

| **Gene** | **Ensembl Identifier** | **No. of Amino acids** | **Predicted Molecular Weight (kDa)** | **Isoelectric point (pI)** | **Sub-cellular Localization** |
| --- | --- | --- | --- | --- | --- |
| *TaVIH1-4A* | TraesCS4A02G283200 | 1055 | 119.52 | 6.29 | Cytosol |
| *TaVIH1-4B* | TraesCS4B02G030400 | 1012 | 114.91 | 6.48 | Cytosol |
| *TaVIH1-4D* | TraesCS4D02G027800 | 1046 | 118.54 | 6.48 | Cytosol |
| *TaVIH2-3A* | TraesCS3A02G319100 | 1036 | 117.78 | 6.10 | Cytosol |
| *TaVIH2-3B* | TraesCS3B02G347000 | 1036 | 117.71 | 5.95 | Cytosol |
| *TaVIH2-3D* | TraesCS3D02G312200 | 1034 | 117.70 | 6.01 | Cytosol |
